# Reducing CRISPR dark matter reveals a strong association between the bacterial membranome and CRISPR-Cas systems

**DOI:** 10.1101/2022.04.26.489349

**Authors:** Alejandro Rubio, Maximilian Sprang, Andrés Garzón, Maria Eugenia Pachón-Ibáñez, Jerónimo Pachón, Miguel A. Andrade-Navarro, Antonio J. Pérez-Pulido

## Abstract

Antimicrobial resistance is widely recognized as a serious global public health problem. To combat this threat, a thorough understanding of bacterial genomes is necessary. The current wide availability of bacterial genomes provides us with an in-depth understanding of the great variability of dispensable genes and their relationship with antimicrobials. Some of these accessory genes are those involved in CRISPR-Cas systems, which are acquired immunity systems that are present in part of bacterial genomes. They prevent viral infections through small DNA fragments called spacers. But the vast majority of these spacers have not yet been associated with the virus they recognize, and this has been named CRISPR dark matter. By analyzing the spacers of tens of thousands of genomes from six bacterial species highly resistant to antibiotics, we have been able to reduce the CRISPR dark matter from 80-90% to as low as 15% in some of the species. In addition, we have observed that, when a genome presents CRISPR-Cas systems, this is accompanied by particular collections of membrane proteins. Our results suggest that when a bacterium presents membrane proteins that make it compete better in its environment, and these proteins are in turn receptors for specific phages, it would be forced to acquire CRISPR-Cas immunity systems to avoid infection by these phages.

## Introduction

The fight against antimicrobial resistant bacteria is one of the major challenges facing mankind in the near future (1). The World Health Organization prioritized a list of multidrug-resistant bacteria in 2017 to support research and development of effective drugs (2). The most relevant species in this list form the so-called ESKAPE group, whose acronym refers to two Gram-positive bacteria (*Enterococcus faecium* and *Staphylococcus aureus*) and four Gram-negative bacteria (*Klebsiella pneumoniae, Acinetobacter baumannii, Pseudomonas aeruginosa*, and *Enterobacter cloacae*). All of them cause infections, some of which are acquired in the hospital environment (3, 4). A major difference between Gram-positive and Gram-negative bacteria, which conditions the way to fight against them, is that Gram-negative bacteria have a second lipid membrane outside the cell wall, which gives them a richer variability of membrane proteins, such as the outer membrane proteins found in this second layer (5).

Bacteriophages can be used to control bacterial growth and are even used to treat infections in humans (6). Thus, bacteria defend themselves against infection by these phages by means of different molecular systems. Restriction-modification systems are by far the most abundant, being present in 83% of prokaryotic genomes, followed by CRISPR-Cas with about 40% (7). The CRISPR-Cas are adaptive immunity systems found in most archaea and in less than half of the bacteria sequenced (8). They provide acquired immune resistance against phages and other foreign nucleic acid molecules such as plasmids, thus restricting gene transfer (9). There are different types of CRISPR-Cas systems based on genes that are part of the different steps of this immune system (adaptation or spacer integration, expression, and interference) and are generically called *cas* (CRISPR-associated genes). ESKAPE bacteria have CRISPR-Cas systems of the most common types, from I to IV, but only in a minimal number of strains, with frequencies ranging from less than 1% to 60% of genomes, depending on the species (10).

The acquired immunity of CRISPR-Cas systems is based on short nucleotide fragments, called spacers, cropped from the foreign nucleic acid sequence of a previous entry into the bacterial cell, called protospacers, which are inserted into the bacterial DNA next to the *cas* genes. These spacers are mostly similar to phage sequences and to a lesser extent to other extrachromosomal nucleic acid molecules. However, a large proportion of them have no known protospacer (over 80-90%). These spacers of unknown origin are believed to originate from as yet unsequenced phages and constitute what has been called the CRISPR “dark matter” (11).

Bacterial genomes with CRISPR-Cas systems have *cas* genes along with the spacers, which are separated by repeats (short identical or nearly identical sequences). However, other genes have been linked with these systems because specific functionalities have been found in strains that have CRISPR-Cas systems. For example, a relationship has been demonstrated with the formation of multicellular structures called biofilms in *P. aeruginosa, Streptococcus mutant* and *Yersinia pestis* (12–14), connections with the regulation of outer membrane proteins have also been described in *Salmonella* Typhi (15), and a specific relationship with virulence have also been shown in multiple bacterial species (16–18).

In earlier work with *A. baumannii*, we found that the group of strains that usually have CRISPR-Cas systems had genes involved in biofilm at a high frequency, and we also found genes encoding proteins with signal peptides and membrane lipoproteins (19). The analysis was based on a pangenome constructed from this species (all of the different genes found in the genomes of the species). The establishment of these pangenomes is now easier due to the large number of genomes available in public databases, and they allow to analyze the accessory genome, which is the set of genes that is not present in all the strains of a species (20) and are usually obtained by horizontal gene transfer (21). If strains with CRISPR-Cas systems have special functions not found in strains without these systems, accessory genes involved in these functions should appear almost exclusively in the former.

We have created a large pangenome for each ESKAPE species and compared strains with or without CRISPR-Cas systems to discover genes present with a significant frequency in the former. Then, these genes have been functionally analyzed and we found that they are enriched in genes encoding membrane proteins, which reveals a possible relationship with phages that infect bacteria and could take advantage of these membrane proteins as entry receptors. In addition, our results demonstrate that the study of thousands of genomes of the same species allows us to reduce the CRISPR dark matter and to trace the origin of most of the spacers found in them.

## Materials and Methods

### Genome collection and annotation

The assembled sequences of ESKAPE species available in the National Center for Biotechnology Information (NCBI) Genome database on June 14, 2021, including complete and draft genomes, were collected (22). Genomes and metadata were downloaded with the tools datasets 12.1.0 and dataformat 12.4.0 (a total of 68,352 genomes). Genomes with a low number of total genes or a low average number of shared genes (>5 times the interquartile range) were removed on suspicion that they did not correspond to the species studied.

The protein-coding genes were predicted using Prokka version 1.14.5 (23), and the pangenome was created by Roary version 3.12.0 with an identity threshold of 90% and the -s parameter for not separating paralogs at this identity threshold (24). Protein sequences were functionally annotated using Sma3s v2 and the UniProt bacterial taxonomic division bacteria 2019_01 as the reference database (25). Gene names provided by Sma3s were preferentially assigned to each protein. When a gene name was repeated, a sequential number separated by two underscores was added. In cases where Sma3s did not assign a gene name, the one proposed by Prokka was taken, if available, preceded by an underscore. A gene was classified as a core gene if it appeared in ≥99% of the genomes of the species.

CRISPR-Cas systems and their specific types were assigned using CRISPRCasTyper 1.4.1 (26). Types I-Fa and I-Fb of *A. baumannii* were distinguished by looking for their different integration site. To discover the spacers of CRISPR-Cas systems, CRISPRCasFinder 4.2.20 was used with default parameters (27). Only CRISPR arrays with an evidence level equal to 4 were considered. Identical spacers were collapsed together, taking into account both chains. The number of sequences of each type for each species are available in Suppl. Table S1.

### Search for specific gene groups in the pangenomes

Antibiotic resistance genes were found by AMRFinderPlus 3.10.1 using the databases of the 6 bacterial species analyzed (28). Virulence genes were found by performing a similarity search with BLASTP 2.9.0+ (29) against the VFDB database version December 2020 (Virulence Factor Database), requiring at least 90% sequence identity and 90% database sequence coverage (30).

Genes encoding membrane proteins were searched for in the functional annotation performed by Sma3s, to which genes encoding outer membrane proteins were specifically added by a BLASTP similarity search against the OMPdb database release 2021, requiring at least 90% sequence identity and 90% database sequence coverage (31).

Genes from plasmids were searched using the annotation ‘Plasmid’ in the UniProt keyword field. Then, genes with ≥90% sequence identity and ≥90% query coverage with a sequence of the PLSDB database v2020_06_23_v2 were added (32). Viral genes were searched following the same protocol but using the IMG/VR database v3 (IMG_VR_2020-10-12_5.1) (33), and viral genes from the functional annotation with Sma3s. Genes that appeared between 2 genes annotated as viral were also added. The Phaster web server was used to search for complete phages (34).

### Search for protospacers

Protospacers, putative genes recognized by spacers, were searched by performing a similarity search with BLASTN and the blastn-short option turned on, using a threshold of ≥95% sequence identity and 100% spacer coverage. Those protospacers that were also found following the same strategy but using sequences from CRISPR repeats instead of the spacers were discarded to avoid misannotated sequences (35).

### Searching for genes associated to CRISPR-Cas types with inference on random forests

We used inference on multiple iterations of random forests to search for genes associated with specific CRISPR-Cas types. For each species, we compiled one dataset containing all strains that have no CRISPR-Cas systems, and other datasets with the strains containing each CRISPR-Cas type present in the data, respectively. From 20 iterations with different random seeds, the most important features were selected and counted. In this context the features are binary indicators of the presence of the genes for each strain. After all iterations, the count of a gene can indicate how often the random forest deemed it important for the difference between the respective CRISPR-containing and CRISPR-deficient genomes. *Cas* genes were removed from consideration, to be able to focus on non-directly related genes. Genes identified as important features in multiple iterations were considered to be associated with CRISPR-Cas systems, when they were more abundant in CRISPR-Cas containing genomes than in non-CRISPR-Cas containing genomes.

The random forest implementation was done in Python with the scikit-learn package 1.0.2 (36). With each iteration, random parts of the datasets were divided into train and test sets with the ratio of 0.8 to 0.2. For all six species and their CRISPR-Cas systems, the random forests achieved average accuracies higher than 0.93 over all iterations. The trained random forest object has the resulting feature importance by the mean decrease in impurity available as a parameter. The permutation feature importance may be more informative for high cardinality features, but since we only have two values for each feature, the mean decrease in impurity feature importance measurement is sufficient.

### Functional enrichment analysis

To discover the functional enrichment of genes associated with specific CRISPR-Cas types, we used the R package TopGO version 2.40.0 (37), which uses GO terms from a specific ontology. The GO terms used were those annotated by Sma3s. Figures were created using the R ggplot2 library in a custom script.

### MLST assignment, gene profiles and molecular phylogenies

MLST numbers were assigned to each genome by compiling the genes used in the species-specific schemes in PubMLST 23 Nov 2021 (38) and searching them in the genome sequences using the mlst program (https://github.com/tseemann/mlst). The MLST number assigned to each genome, along with the CRISPR-Cas systems it has is available in Suppl. Table S3.

MLST phylogenetic trees were constructed using the MLST sequences. Nucleotide sequences were aligned with mafft v7.271 using the G-INS-I option (39). The phylogeny was constructed with RAxML v8.2.9 with the GTRCAT model and bootstrap of 1000 (40). The model was selected with ModelFinder implemented in IQ-TREE (41). The phylogeny in Fig. 6d was constructed with the same protocol but using all genomes, the PROTGAMMAWAG model and 500 core proteins.

The gene profiles for each species were constructed using a binary representation of the bacterial genome, where a gene is either absent or present in a strain, without accounting for the number of paralogs. This data is condensed to MLST level, by assigning 0 or 1 to the gene in the MLST group by majority vote of all strains in the group. The MLST groups are then subjected to a pairwise Jaccard distance measurement, resulting in a N x N matrix of Jaccard distances between MLST groups, with N equal to the number of MLST groups for the respective species dataset. The pairwise Jaccard distances were computed with scikit-learn.

These pairwise distances were used to construct a profile of genetic distances between the MLST groups for each species. We used ward-linkage and descending distance sort for the hierarchal clustering and the dendrogram. Dendrograms were produced using SciPy 1.6.2 (42), correlation plots were plotted with seaborn 0.11.2 and matplotlib 3.5.0 (43, 44).

## Results

### A large proportion of CRISPR dark matter spacers could be annotated by the pangenome analysis

The genomes of the different ESKAPE species were initially obtained and they were both structurally and functionally annotated with a special emphasis on the protein-coding genes and the elements that are part of the CRISPR-Cas systems. According to the number of genes, the smallest pangenome was found for *S. aureus* despite having started from a larger number of genomes (Fig. 1ab). The species with the largest number of genomes with CRISPR-Cas systems were *K. pneumoniae* and *P. aeruginosa*, which also had the largest pangenomes (along with the other Gram-negative species) and more than 3500 core genes, i.e., genes common to all genomes of the species. The difference between Gram-negative and Gram-positive bacteria is also evident when comparing the number of genes per genome, with *P. aeruginosa* showing both the largest number of genes and average number of shared genes (Fig. 1c). Finally, the major difference between genes per genome and average number of shared genes is found in *E. cloacae*, which could be explained by the low number of genomes used for this species (n=317).

**Fig. 1.**
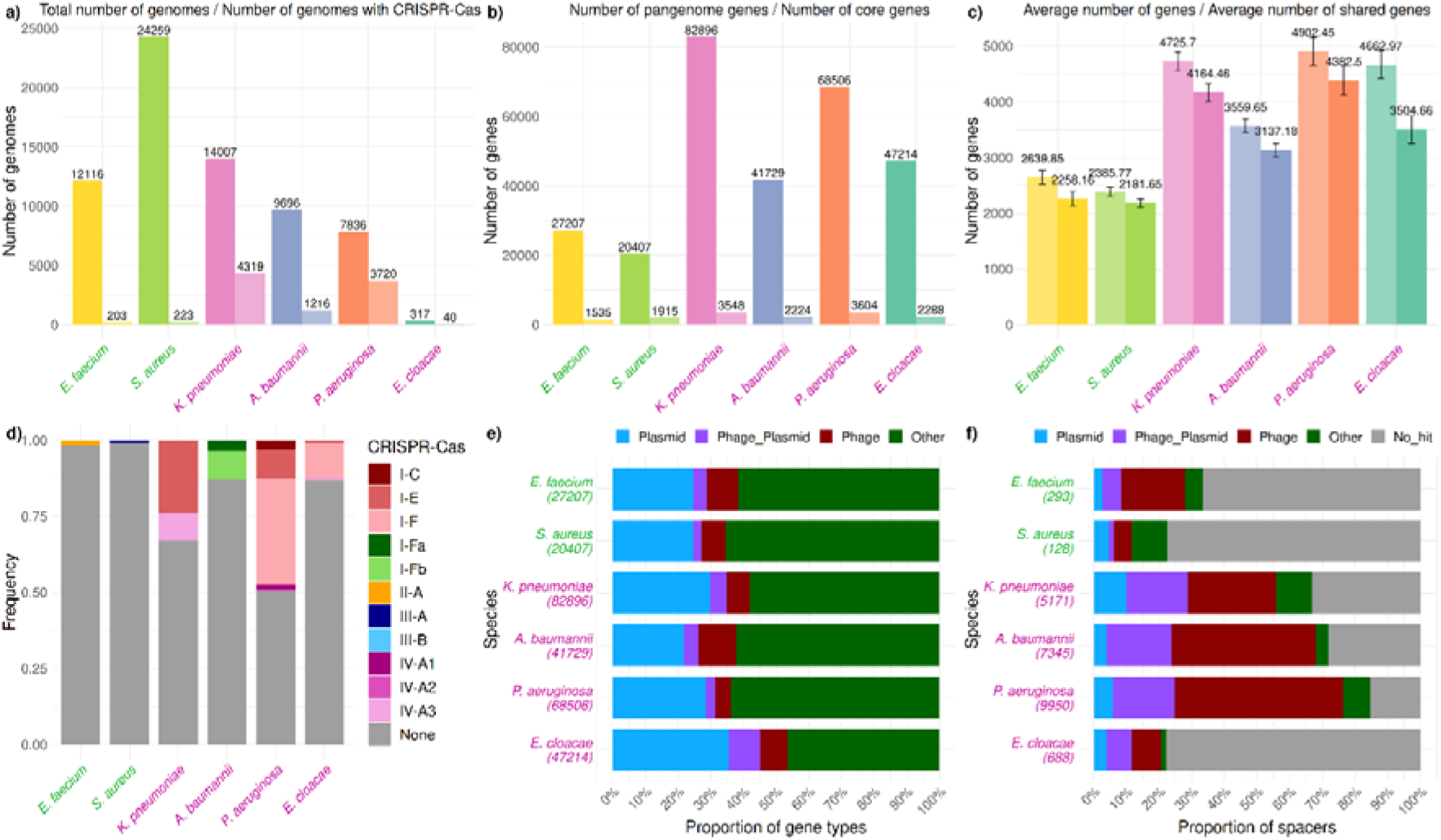
Summary of analyzed genomes. a) Total number of genomes analyzed and number of genomes having CRISPR-Cas systems. b) Number of genes in the pangenome and number of core genes. c) Average number of genes per genome and average number of shared genes (average of the number of genes shared for each genome with the remaining genomes). d) Distribution of CRISPR-Cas systems among genomes. e) Proportion of gene types in the pangenome. For each species: the number of total genes is shown (below the species name), and the proportion of them that were annotated as belonging to plasmid sequences, phages, both plasmids and phages, and other genes not included in plasmids or phages. f) Spacers and assigned protospacer types by species. For each species: the number of total spacers is shown (below the species name), and the proportion of them that match with plasmid sequences, phages, sequences annotated as both plasmid and phage, other genes not included in plasmids or phages, and the spacers that do not match any gene (No_hit). The names of Gram-positive bacteria appear in green, and those of Gram-negative bacteria in purple.

The overall proportion of genomes exhibiting CRISPR-Cas systems is low, with Gram-positive bacteria having them in only 1% of their genomes, both *A. baumannii* and *E. cloacae* in 12%, and *K. pneumoniae* and *P. aeruginosa* in 30% and 47%, respectively (Fig. 1d). Since the spacers of CRISPR-Cas systems usually recognize mobile genetic elements, we first separated pangenome genes that could come from plasmids (an average of 28% of the genes) and bacteriophages (an average of 8% of the genes) (Fig. 1e). Interestingly, an average of 5% of the genes were annotated as originating from both genetic elements, which could come from what is known as phage-plasmids (45).

Next, the spacers were obtained and their cognate protospacers were searched for within the pangenome genes. The major number of different spacers was found in the three species with CRISPR-Cas IV and/or I-F types (Fig. 1f). As expected, protospacers belonged in a much larger proportion to phage genes. Remarkably, it was possible to annotate more than 65% of the spacers in the same three species mentioned, with a maximum of 85% in *P. aeruginosa*. The species with a lower number of annotated spacers (about 25%) were *E. cloacae* and *S. aureus*. It is noteworthy that about 18% of the spacers in CRISPR-Cas type I species appear to recognize genes annotated as phage-plasmids.

Since the presence of CRISPR-Cas systems prevents the entry of foreign DNA (including resistance and virulence plasmids) into the bacterium, the number of genes involved in these functions was compared between genomes with and without CRISPR-Cas systems. On average, genomes with CRISPR-Cas systems presented a lower number of resistance and virulence genes (Suppl. Fig. S1). The most significant difference was found with CRISPR-Cas type II and III, except for resistance genes in *S. aureus*. On the other hand, *P. aeruginosa* presented the highest number of virulence genes overall, although the genomes without CRISPR-Cas systems presented a lower number than those with CRISPR-Cas systems, a result that was repeated with the resistance genes. So, we cannot conclude that all genomes with CRISPR-Cas systems tend to carry a lower number of resistance and virulence genes.

### CRISPR-Cas systems appear and disappear throughout the entire phylogeny

At this point we wanted to know if the CRISPR-Cas systems were linked to a branch of the phylogeny of the species studied. To define phylogenetic relationships between genomes with and without CRISPR-Cas systems, the multi-locus sequence typing was used (MLST). This is based on several house-keeping genes of each species (38), and two adjacent groups reflect genomes arising from a recent common ancestor. Except for *E. faecium* type II and other specific aggregations in other species, CRISPR-Cas systems appear to be spread throughout the phylogenetic tree (Fig. 2, top), suggesting a possible gain by horizontal gene transfer in genomes for which it could provide an evolutionary advantage, and a possible subsequent loss when that advantage no longer exists.

**Fig. 2.**
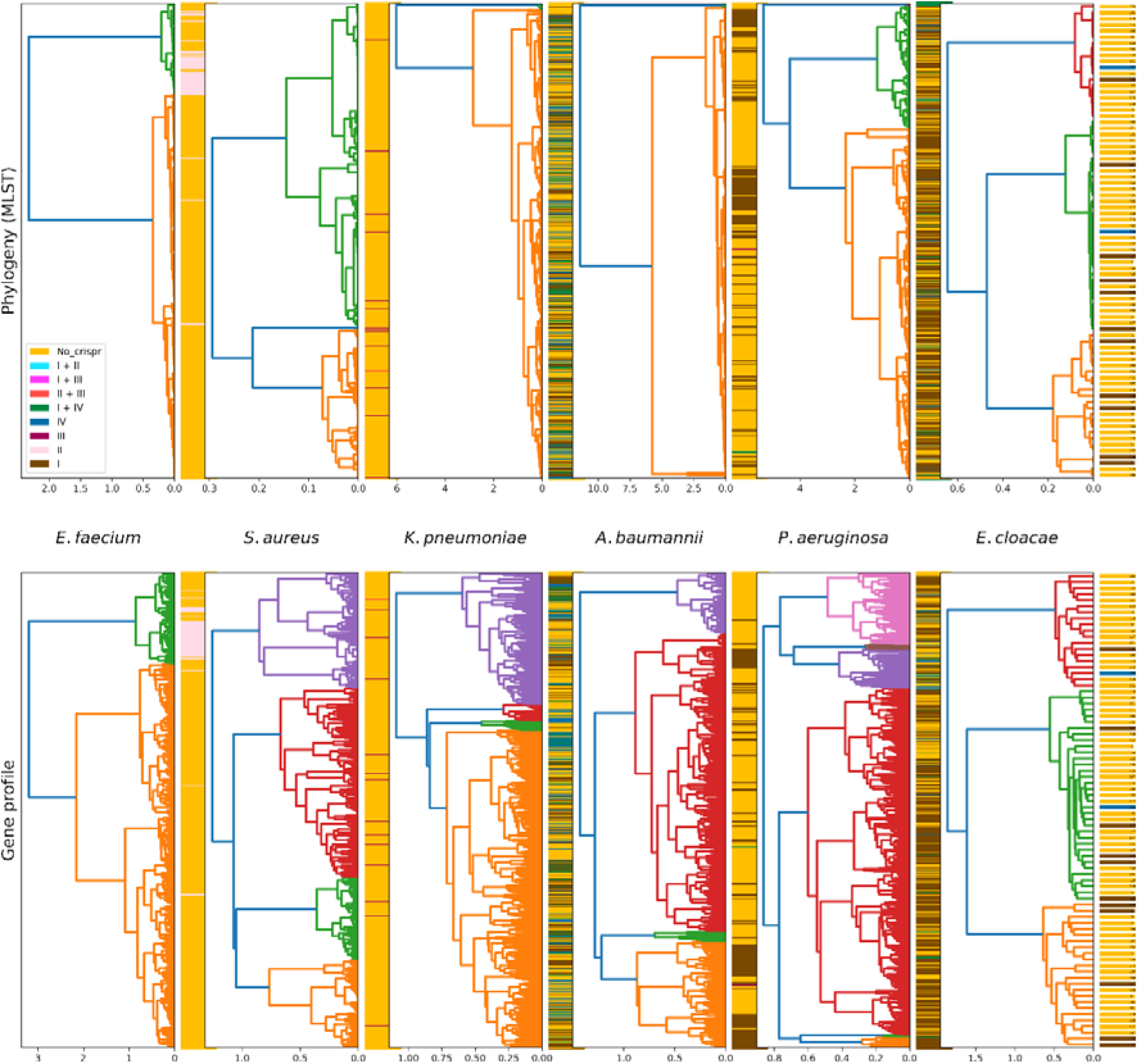
Molecular phylogeny for each species based on MLST (top) and gene profile (bottom). The legend shows the different types of CRISPR-Cas (without specifying the subtype) with orange indicating the MLST group lacking any CRISPR-Cas system. The colors of the branches highlight different clades distanced from the rest.

If CRISPR-Cas systems are associated with other bacterial physiological functions, one possible hypothesis is that genomes with CRISPR-Cas systems always have a similar collection of accessory genes, regardless of *cas* genes. To test this idea, distance trees were constructed between the same MLST groups in the phylogeny, in this case based on the gene profile of the genomes (gene presence/absence matrix). These gene profiles showed a similar dispersion to that found with molecular phylogeny, except, again, for some particular aggregations, suggesting that CRISPR-Cas systems do not appear in genomes with a fixed collection of accessory genes (Fig. 2, bottom). However, when the two types of relationships are compared, a high correlation is observed between distances of the molecular phylogeny and the gene profile in species with a smaller number of genomes with CRISPR-Cas systems (Suppl. Fig. S2). Nevertheless, this correlation decreases in those with a higher number of genomes with CRISPR-Cas systems. Taken together, this reinforces the idea that CRISPR-Cas systems do not appear to be linked to strains phylogenetically related or to specific accessory genomes. But this raised the question of whether genomes with CRISPR-Cas systems presented particular genes at higher frequencies than genomes without these systems.

### Genes associated with CRISPR-Cas systems mainly encode membrane proteins

We have already seen that CRISPR-Cas systems are associated with different accessory genomes and appear and disappear at any evolutionary time. However, assuming that CRISPR-Cas systems are associated with other physiological functions of bacteria, we should find some accessory genes more frequently in genomes having these systems. With this in mind, we searched for genes significantly associated with genomes showing CRISPR-Cas systems, excluding the *cas* genes themselves. Thus, a mean of 147±62 genes per genome were found associated to the different CRISPR-Cas systems, while only 39±47 genes were associated with not having CRISPR-Cas systems (Suppl. Table S2). To test whether the CRISPR-Cas associated genes were significantly involved in any biological process or function, enrichment analysis was performed, and it was found that genes encoding membrane proteins were highly prominent (Fig. 3). These included the pili proteins of *P. aeruginosa* and *K. pneumoniae*, as well as outer membrane proteins of *A. baumannii* and *K. pneumoniae*, and other *A. baumannii* and *P. aeruginosa* proteins involved in type II secretion systems. This relationship with membrane proteins was especially relevant in CRISPR-Cas type I systems, while other genes involved in catabolic processes or DNA metabolism were found more notably in types II, III, and IV (Suppl. Fig. S3). In addition, types IV were also notable for the annotation “extrachromosomal DNA”, reflecting the origin of these systems from mobile genetic elements (46).

**Fig. 3.**
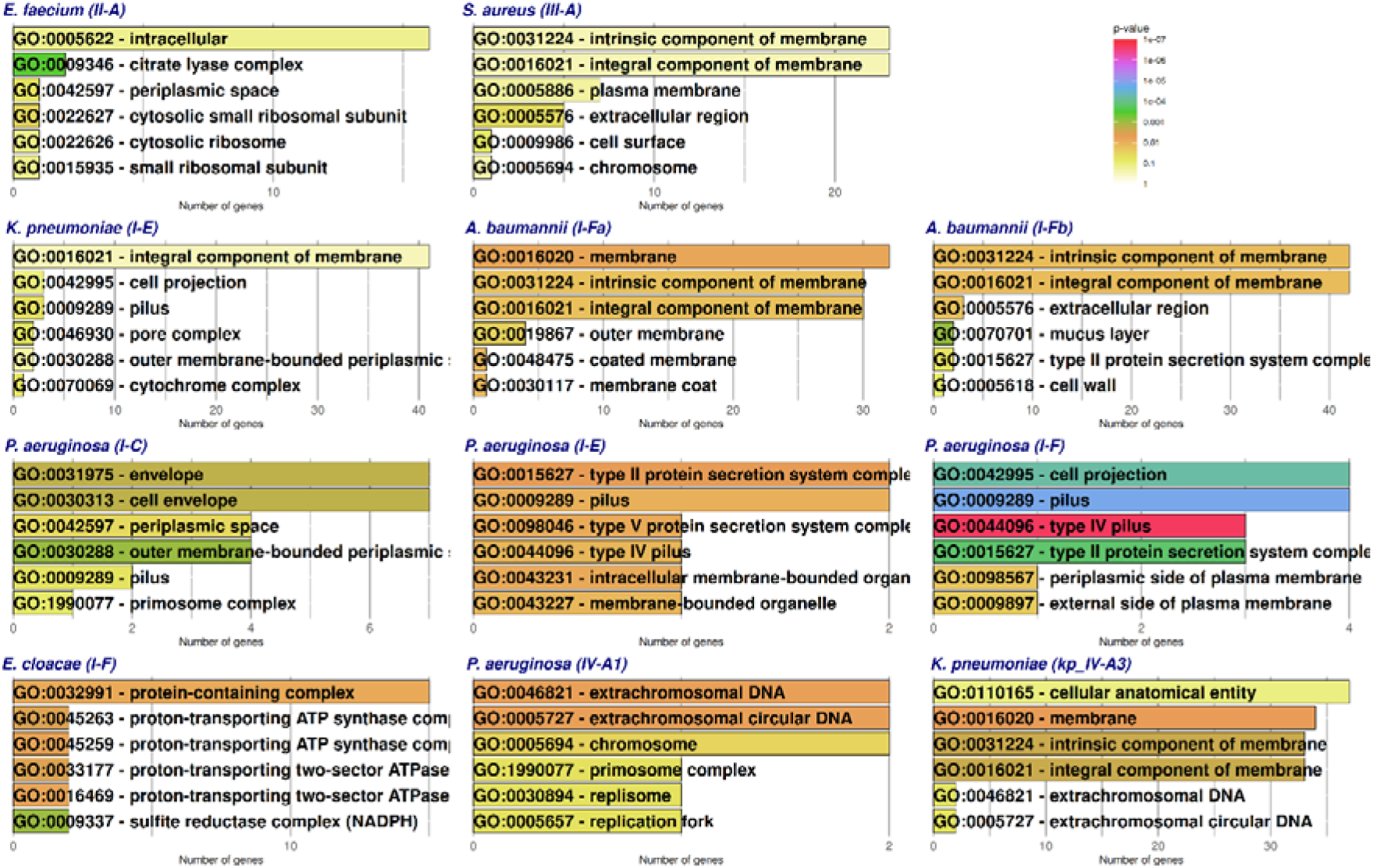
Functional enrichment of genes associated with different CRISPR-Cas systems in the different species. Gene Ontology cellular component was used in these enrichments, using functional annotations of the pangenome obtained by Sma3s. Enrichment with Gene Ontology biological process and molecular function can be found in Suppl. Fig. S3.

### Genomes with specific types of CRISPR-Cas systems have different sets of membrane proteins

Since the genomes bearing CRISPR-Cas systems presented specific types of membrane proteins, we wanted to know if all genomes with a specific CRISPR-Cas type had the same set of membrane proteins. To assess this, genes encoding membrane proteins previously associated with CRISPR-Cas were searched for in genomes having these systems. In general, there were at least two clusters of genomes, especially when type I was analyzed, each one presenting a different collection of genes encoding membrane proteins (Fig. 4), except for type I-F in *E. cloacae*, as the number of genomes in this species is low. However, the separation between these two clusters of genomes were not as clear as with the other types (Suppl. Fig. S4). As an example, about half of the *A. baumannii* genomes with the CRISPR-Cas type I-Fb system carry the *opuD* and *betP* genes, involved in choline and glycine betaine transport, which could protect the bacterium from osmotic stress (47). And the other half of the genomes have porins such as *benP*, or a specific variant of the TonB-dependent siderophore receptor *bauA* (48). The clusters found with CRISPR-Cas systems genomes do not seem to be normally associated with the phylogeny of the corresponding species, since the genomes that present the membrane proteins that define them are distributed throughout this phylogeny (Suppl. Fig. S5).

**Fig. 4.**
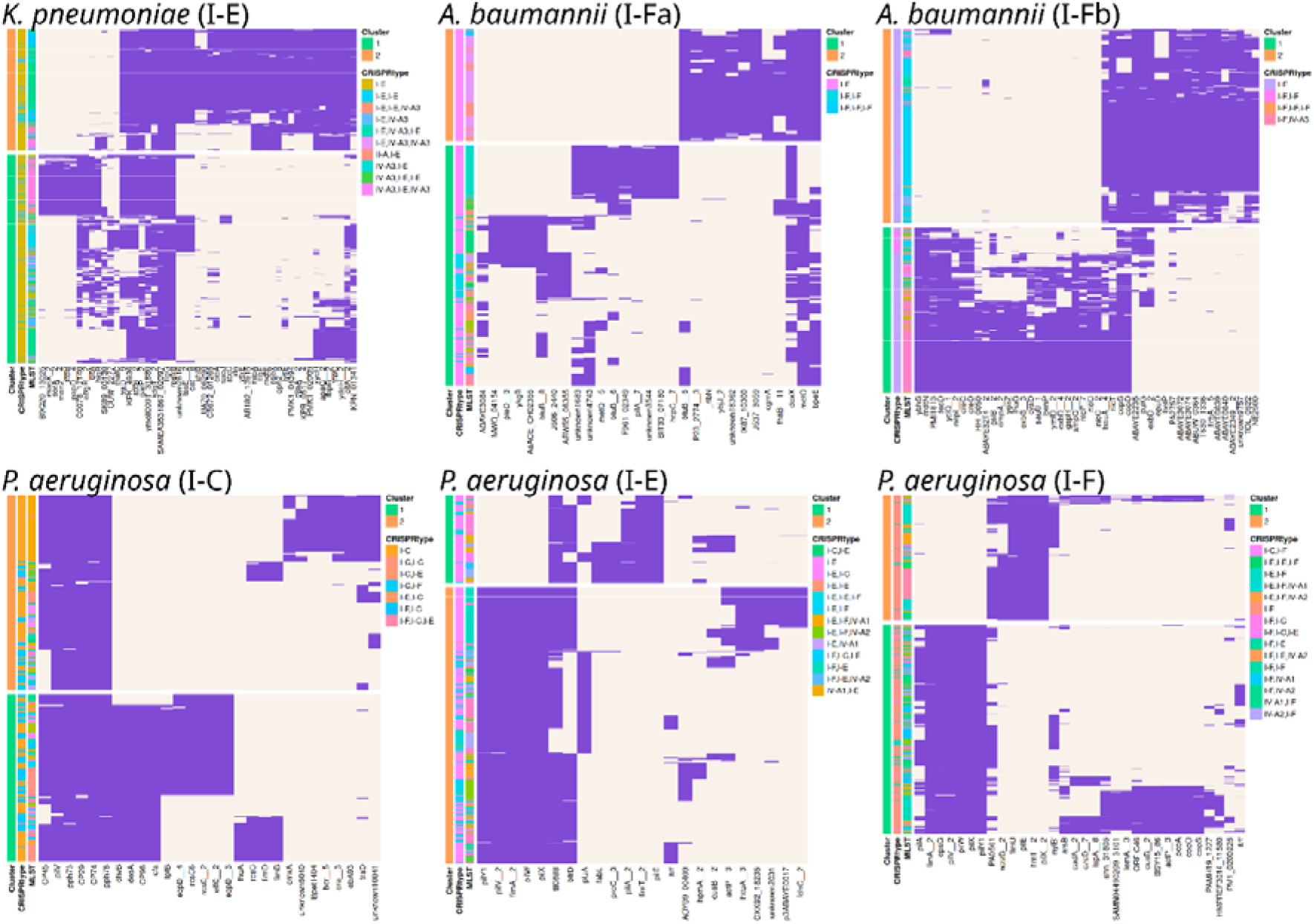
Clusters of genomes with CRISPR-Cas type I systems according to the relevant genes encoding membrane proteins that they have. The purple color indicates presence of the gene (X-axis) in the corresponding genome (Y-axis). On the left side of each plot, the cluster number is shown, along with the CRISPR-Cas type or combination of them and the MLST group.

### Genomes with both CRISPR-Cas type I systems and specific membrane proteins show exclusive spacers and phage genes

Genomes with CRISPR-Cas type I systems show specific collections of membrane proteins. These proteins could provide the bacterium with important characteristics such as specific stress responses, or the detoxification that certain outer membrane proteins can offer. Since the majority of CRISPR-Cas systems studied here seem to recognize phage sequences (Fig. 1f), we hypothesized that these membrane proteins could be acting as receptors of specific phages, and genomes with them are forced to acquire CRISPR-Cas systems to defend against infection by these viruses, while maintaining the beneficial functions for the bacterium. Indeed, the *A. baumannii* cluster 1 for type I-Fb includes the *ompA* gene, an outer membrane protein, the *P. aeruginosa* cluster 1 for type I-C shows the *fhuA* gene and the two *A. baumannii* clusters for I-Fa show different variants of the *btuB* gene, both encoding TonB-dependent proteins. These three genes have long been known to act as phage receptors in *Escherichia coli* and *Salmonella* (49).

To test the above hypothesis, we looked for spacers that recognized phage genes present in each of the two membrane protein-based clusters of genomes that did not appear (neither the spacer nor the phage gene) in the other cluster. We then searched for genomes in each cluster that might carry the phage genes recognized by these spacers. Thus, within a given cluster we could find genomes containing a cluster-specific spacer, or a phage gene recognized by one of these spacers. But it could also be the case that both elements appear in the same genome, and this would imply having two new alternatives: either a cluster-specific spacer and the viral gene recognized together, or a spacer and a viral gene recognized by one of the other cluster-specific spacers. When evaluated in CRISPR-Cas type I systems, the number of genomes with unique spacers for each cluster was high, with a predominance of genomes with unique spacers in *K. pneumoniae* I-E and *P. aeruginosa* I-C, with spacers and phage genes in *A. baumannii* I-F types, and with phage genes in *P. aeruginosa* I-E and I-F (Fig. 5).

**Fig. 5.**
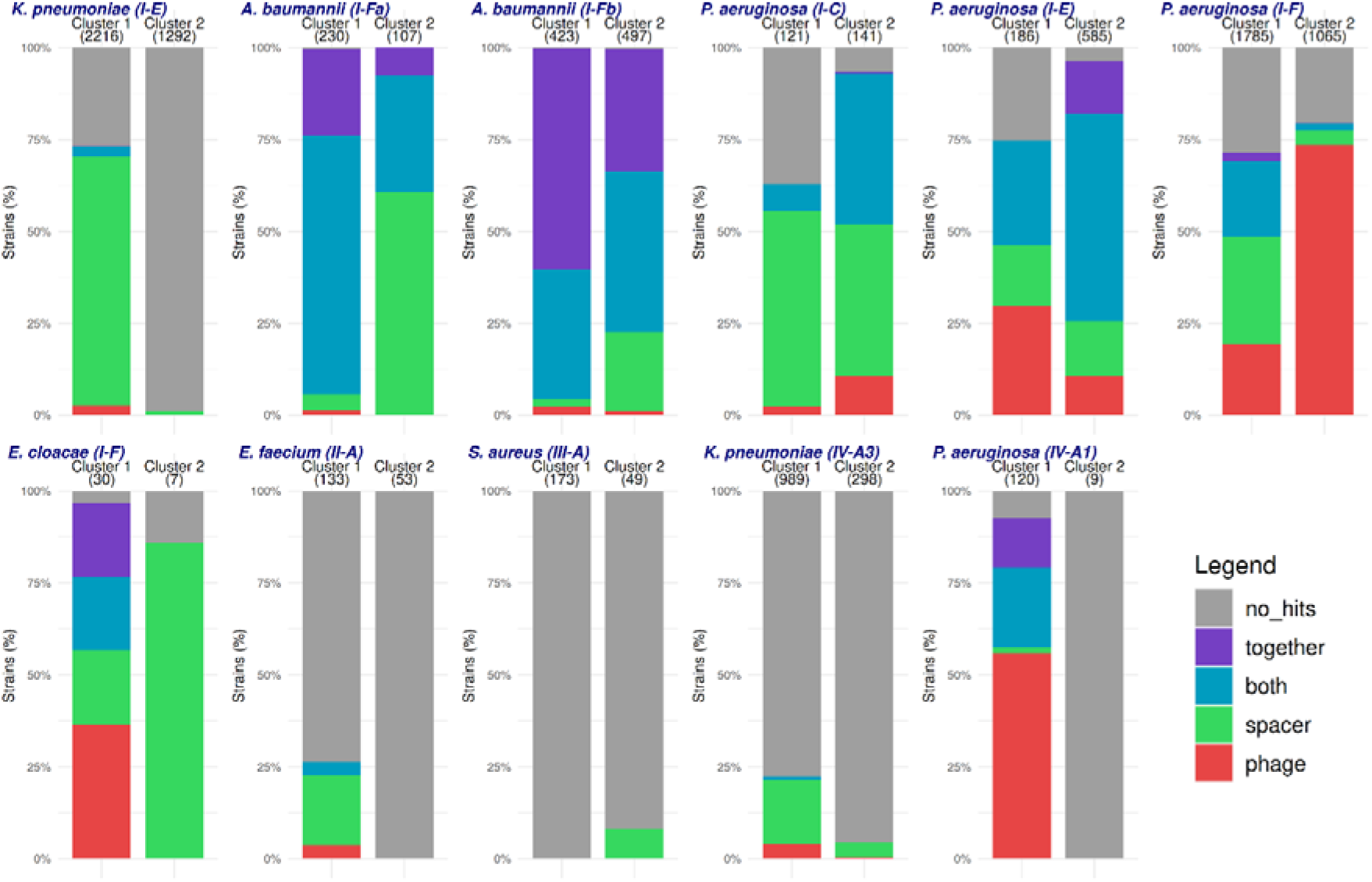
Distribution of genomes in each membrane protein-based cluster with unique spacers and/or protospacers (phage genes). The chart shows the proportion of genomes from each cluster which contain: a phage gene recognized by a cluster-specific spacer (red), a cluster-specific spacer (green), both a spacer and a phage gene, but the latter is recognized by another cluster-specific spacer (blue), both a spacer and the recognized phage gene (purple), or neither the cluster-specific spacer nor the phage gene (grey). The types of CRISPR-Cas systems that defined the best clusters are shown in the row above (in the same order as in Fig. 4), and the others are shown in the row below (in the same order as in Suppl. Fig. S4). The number of genomes in each cluster is shown in parentheses.

In *A. baumannii*, cluster 2 of CRISPR-Cas I-Fb genomes (see Fig. 5) has specific spacers against a phage gene (*unknow5433*) that appears in 466 distinct genomes, with the phage gene itself also appearing in 68 of them (Fig. 6a). The complete phage is integrated near a tRNA-Arg gene and has a total of 59 proteins (Suppl. Fig. S6). In addition, the phage gene also appears in 283 genomes lacking CRISPR-Cas systems. When the specific membrane proteins of cluster 2 are also searched for in genomes lacking the CRISPR-Cas system but having the phage gene, the best match occurs with the *T630_1336* membrane protein (which we will refer to as *cam1* for CRISPR-associated membranome gene 1), while the rest of the membrane proteins appear more frequently in genomes lacking the phage gene (Fig. 6b). Specifically, 278 of the genomes that have *cam1* also have the phage gene (63%). In fact, of the 283 genomes without CRISPR-Cas systems that have the phage gene, only in 5 of them the *cam1* gene could not be found.

Cam1 is a 349 amino acid protein that shows a transmembrane region followed by a Rhs repeat-associated core domain at its N-terminal half (InterPro:IPR022385). This domain appears in bacterial toxins involved in type VI secretion systems (T6SS), but when present in proteins of less than 400 amino acids, found in bacteria such as *Pseudomonas putida*, it has an unknown function (50). The gene encoding this membrane protein is part of a cluster of 6 genes that is integrated next to a tRNA-Glu and the gene *fhuA*, and close to this region there is another T6SS spike gene (*vgrG2__8*). Since the presence of this protein seems to be a *sine qua non* condition for the appearance of the phage, we could expect that the gain of this putative T6SS by the bacterium would imply that it would be exposed to infection by the phage, something that could be counteracted by the bacterium with the acquisition of a CRISPR-Cas system (Fig. 6c). This appears to be supported when analyzing the group of genomes containing this membrane protein, together with genomes that have a similar gene profile. There is a set of 65 genomes with a gene profile similar to cluster 2 CRISPR-Cas I-Fb genomes which lack the membrane protein, phage and CRISPR-Cas systems (Fig. 6d). Then, the phylogeny of this group of genomes shows a first divergence supporting the gain of the membrane protein Cam1 that seems to allow phage entry. Later in the phylogeny, a new divergence allows the gain of the CRISPR-Cas system that seems to prevent phage integration. Indeed, while genomes with only the phage gene or CRISPR-Cas systems are rare (5 and 21 genomes, respectively), there are 278 genomes with both Cam1 and the phage gene, and 401 with both Cam1 and the CRISPR-Cas system but not the phage gene (Fig. 6e), supporting the dependence of phage on Cam1, and the dependence on CRISPR-Cas systems to prevent phage.

**Fig. 6.**
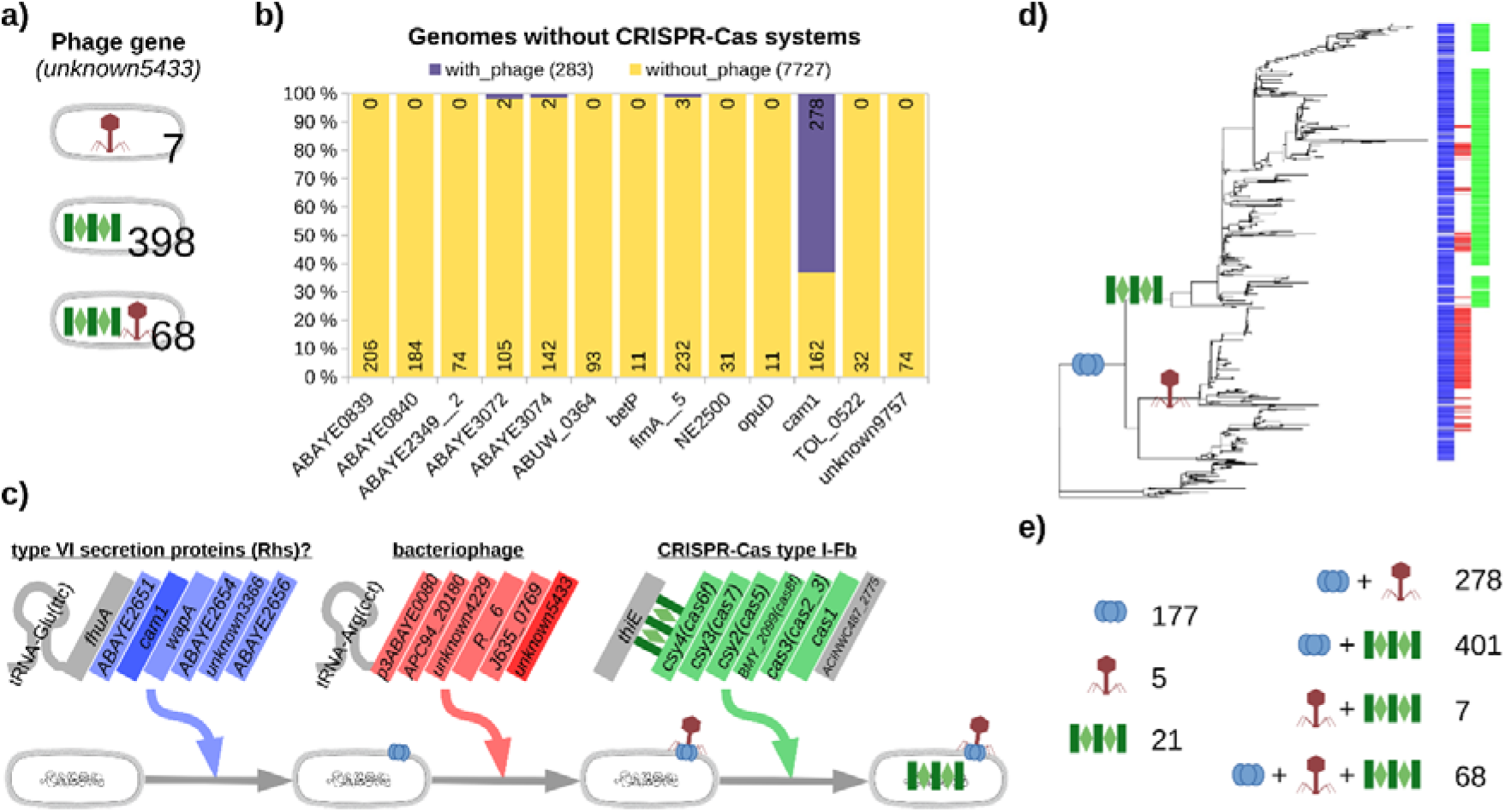
Phage gene specific to cluster 2 of CRISPR-Cas I-Fb genomes and its co-occurrence with membrane genes. a) The phage gene *unknown5433* appears alone in only 7 genomes of cluster 2, but there are spacers against it in 398 genomes, and the phage gene appears together with these spacers in 68 other genomes. b) Frequency of cluster 2-specific membrane proteins in genomes lacking CRISPR-Cas systems, separated between those with and without the phage gene. The number of genomes with and without the phage gene is shown in parentheses. Note that the phage gene appears in most genomes having the membrane protein Cam1. c) Hypothesis of gain of gene groups from a genome lacking the membrane protein and the CRISPR-Cas system: first the region of 6 genes involved in a T6SS entered, which would allow virus entry through the Cam1 protein, and finally the CRISPR-Cas system would be obtained for protection against the phage. d) Molecular phylogeny of 727 genomes of *A. baumannii* that belong to the membrane protein-based cluster 2 or have a similar gene profile to genomes in this cluster. Five hundred genes that appeared in all genomes were used. The metadata columns highlight the genomes with the membrane protein Cam1 (blue color), the phage gene *unknown5433* (red color), and the CRISPR-Cas type I-Fb (green color). A genome of the reference MLST2 group was used to root the tree (Strain XH727). This tree would be consistent with the steps proposed in the hypothesis in c). e) The number of total genomes of *A. baumannii* possessing each possible combination of elements (Cam1, phage gene and CRISPR-Cas I-Fb).

## Discussion

A limited number of bacterial genomes have CRISPR-Cas systems to prevent the entry of foreign DNA. We have analyzed tens of thousands of genomes of bacterial species of the ESKAPE group and found that type I is the most frequent CRISPR-Cas system in the Gram-negative species of this group. Our results are consistent with previous reports that found, for example, a low proportion of genomes with CRISPR-Cas systems in *E. faecium* (51), and about fifty percent of those in *P. aeruginosa* (9). When a bacterial genome has a CRISPR-Cas system, it is expected to have fewer genes. Some of the missing genes may be those involved in antibiotic resistance and originating from plasmids or integrative and conjugative elements, or those involved in virulence and originating from phages. This was mostly found in our study with ESKAPE pangenomes, and coincides with previous studies with the same species, except for the fact that we show a more complete collection of type IV systems, and we have used twice as many genomes (10, 52).

Bacteria have different defense systems against phages. But to study *in silico* the relationship of these systems with their target sequences, CRISPR-Cas systems have the advantage of having spacers, which reflect previous encounters with exogenous sequences (7). Thus, spacers of CRISPR-Cas systems usually recognize sequences originating from phages (11) and to a lesser extent from plasmids, such as we previously showed in *A. baumannii* (19). However, most spacers have unknown origin, as the corresponding protospacer cannot be found, and have been lumped into what is known as CRISPR dark matter (11, 53). This dark matter accounts for 80-90% of the spacers and is expected to recognize phage sequences that are still unknown or have diverged from known phage sequences. And it is believed that this percentage may decrease with the future increase of sequences in the databases (54). We show that by analyzing complete pangenomes, which can include all the variation of phages infecting the species, dark matter can be greatly minimized (Fig. 1f). Thus, we were able to annotate 85% of the 9950 *P. aeruginosa* spacers and 72% of the 7345 *A. baumannii* spacers, with approximately 70% of these corresponding to phages or phage-plasmids. These phage-plasmids include phages that can remain as extrachromosomal elements in the bacterium (45) and against which we have found more associated spacers than against plasmid genes.

We have also found that genomes with CRISPR-Cas systems do not appear phylogenetically restricted, nor do they have a unique accessory genome. This suggests that bacteria acquire these systems when they provide an important evolutionary advantage. In a study carried out to measure the impact of CRISPR-Cas systems on horizontal gene transfer in bacteria, it was concluded that these systems would play an important role at the population level but not at the evolutionary scale (55). In fact, these systems are often recruited by mobile genetic elements on independent phylogenetic times (56). All of this would support our results, positioning CRISPR-Cas systems as functional modules that are acquired and discarded under certain circumstances.

We found dozens of genes that co-occur with CRISPR-Cas systems, although not necessarily close in the bacterial chromosome, many of which encode membrane proteins. This association suggests that one such circumstance could be the defense against phages that use these proteins as receptors. A previous report had already suggested that misfolded membrane proteins may trigger an envelope stress response that activates a CRISPR-Cas system (57), and other reports have found genes with probable association to the CRISPR-Cas systems and some of them encoded integral membrane proteins (58, 59). Many of these genes were related to type III systems, which is the type in *S. aureus*, where we found more than 20 genes annotated as integral component of membrane associated with its CRISPR-Cas system. However, we mainly found this association with membrane-related accessory genes among the different classes of type I CRISPR-Cas and propose that this may be related to the acquisition of beneficial functions for the bacterium that conversely make it more vulnerable to certain phages. These membrane proteins may help form biofilms or allow for certain virulence-related advantages (12–18), but at the same time, this membrane proteins can be receptors for specific phages.

It has been shown that phages can increase the virulence of the bacterium they infect when integrated into the bacterial chromosome, as they can carry toxins, resistance genes, or adhesion factors (60). CRISPR-Cas systems would prevent these phages from proliferating. However, we have found that in many cases spacers coexist with the cognate phage gene, especially in *A. baumannii*. These cases reflect that the phage would be integrated into the bacterial genome, suggesting that the immune system has not been fully efficient. Indeed, in *P. aeruginosa*, and partly in *K. pneumoniae*, we have seen that the number of virulence genes in strains carrying CRISPR-Cas systems may be higher than in those without (suppl. Fig. S1). The coexistence of the protospacer with the cognate spacer has been proposed as representing autoimmunity processes with a negative effect on the bacterium (61), although this may also be explained by the fact that the prophage is expressing anti-CRISPR systems (62). However, other studies have shown that CRISPR-Cas systems can prevent the lytic cycle of phages but tolerate the virus integration as a prophage, allowing the bacteria to co-opt the phage genes for possible use as virulence factors (63).

We observed that genomes with specific types of CRISPR-Cas systems can be separated into clusters based on the membrane proteins they have, and that the spacers of these CRISPR loci would recognize different non-overlapping phage genes (Fig. 4-5). By analyzing these relationships between membrane proteins, spacers and phage genes, we are able to propose a gene encoding a member of a putative T6SS as a putative receptor for a phage found in genomes with and without CRISPR-Cas systems. Phylogenetic data suggest that the gain of this immune system would protect the bacterium against this phage while allowing it to maintain the secretion system, which could be useful for intra- or interspecific competition (64). Interestingly, the gene cluster to which this membrane gene belongs is integrated next to the *fhuA* gene, and it is speculated that a class of specific receptors (TonB) could be critical for phage genome injection through their interaction with FhuA (65). Thus, this protein could help both the entry of the phage and the proper functioning of the T6SS system. Other reports have shown that secretion systems and CRISPR-Cas systems can depend on quorum sensing (66, 67), which enables the coordination of bacterial population growth and is therefore also related to biofilm formation. Thus, it could be hypothesized that the bacterium would express both systems at the same time, to avoid being exposed to phages that could take advantage of the activation of the T6SS system when the bacterial population and cell-to-cell contact is increased.

## Conclusions

We demonstrate that the use of large pangenomes allows to annotate a great part of the spacers of CRISPR-Cas systems, which will allow further research in this field. Here, we describe a “Membrane protein-Phage-CRISPR” triad, in which the CRISPR-Cas systems might be especially necessary when the bacterium expresses accessory genes that encode for membrane proteins. This would give it a special advantage in that situation, but it would also represent a gateway for phages that use those proteins as receptors.

## Data availability

All data generated or analyzed during this study are included in this published article and its supplementary information files.

The code used to analyze the data and the pangenomes as well as the phylogenetic distances can be found in our GitHub repository: https://github.com/UPOBioinfo/crispromeskape.

## Acknowledgements

We would like to thank C3UPO and the HPC-group of the JGU for the HPC support.

## Supplementary data

**Suppl. Fig. S1.**
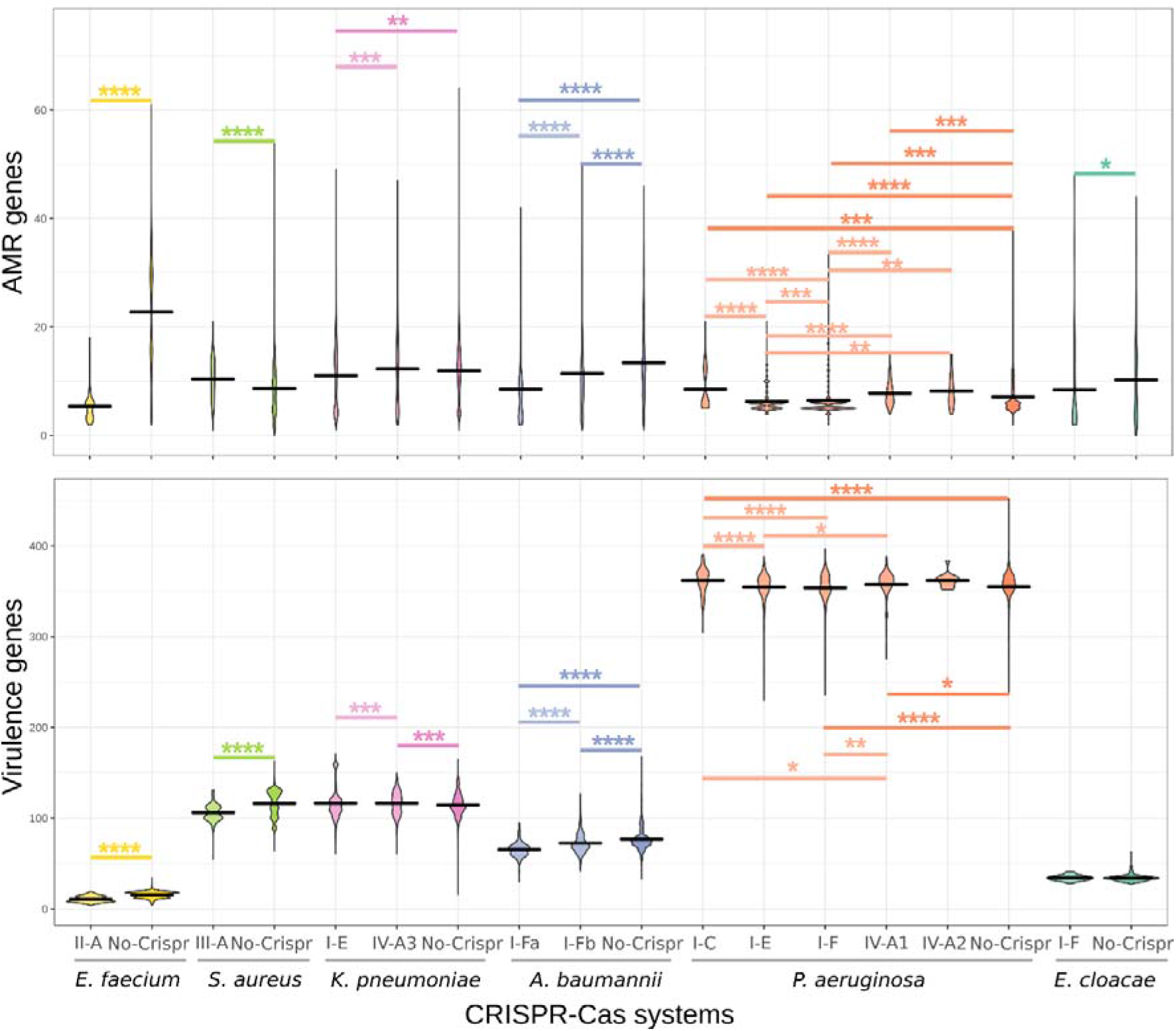
Distribution of the number of genes involved in antibiotic resistance (top) and virulence (bottom) in groups of genomes with and without CRISPR-Cas systems. Asterisks highlight pairs of groups with significant differences in the median according to a Wilcoxon rank-sum test (from *≤0.05 to ****≤1e-03).

**Suppl. Fig. S2.**
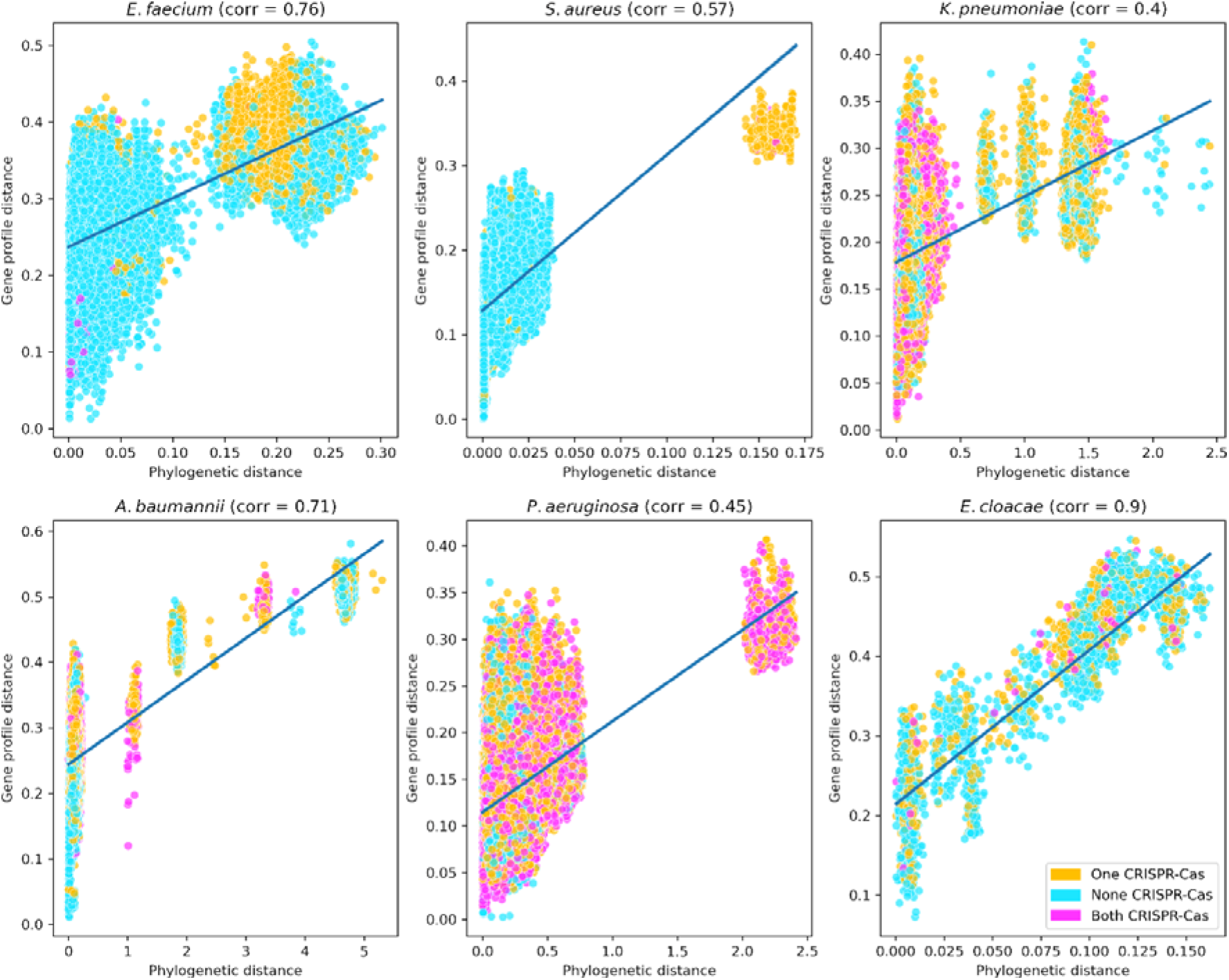
Correlation between molecular phylogeny and gene profile. Pearson correlation value is shown in parenthesis. Points show pairs of MLST groups without CRISPR-Cas systems (cyan), with one group having one of these systems (orange), and with the two groups having CRISPR-Cas systems (pink).

**Suppl. Fig. S3.**
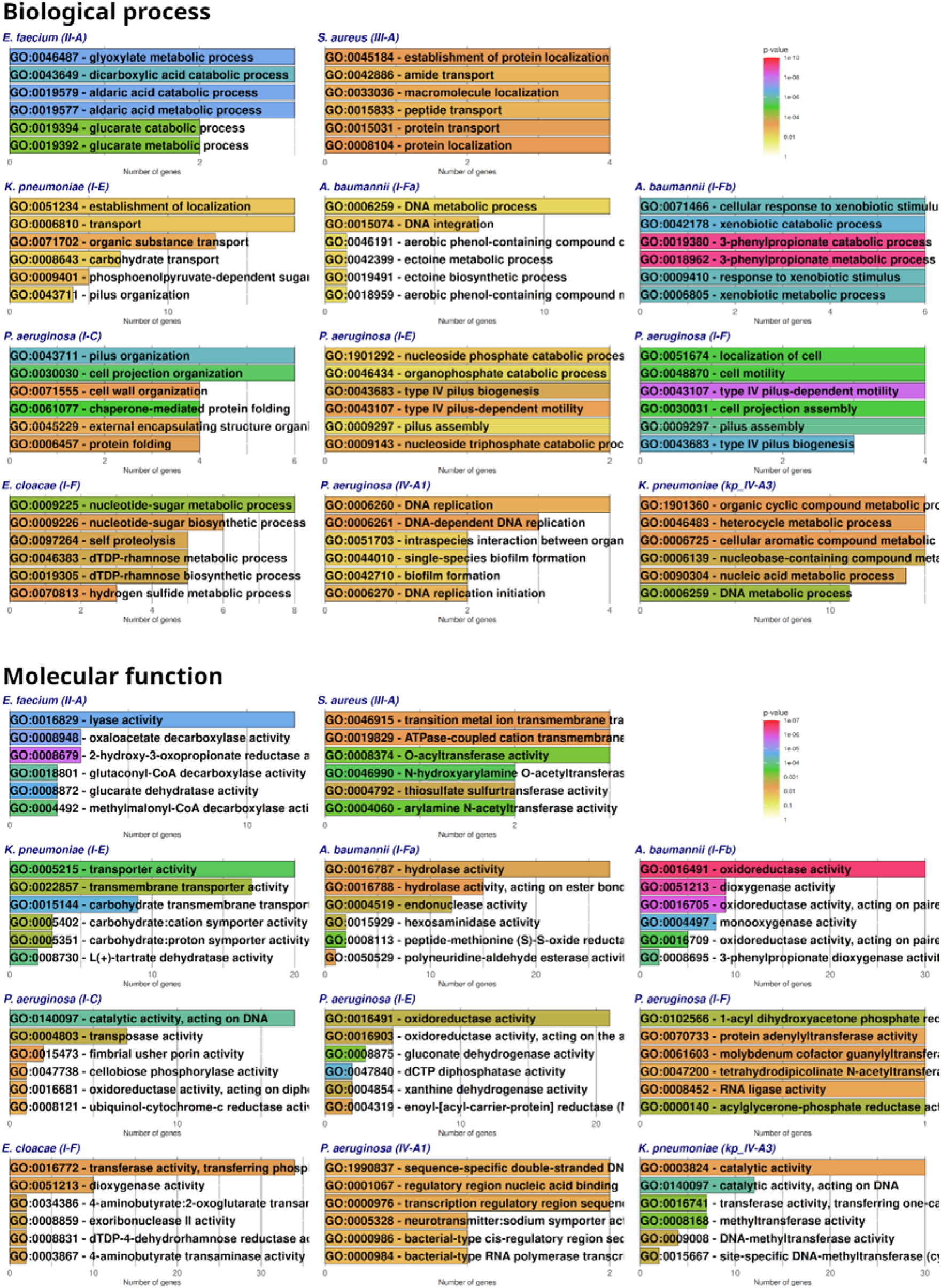
Functional enrichment of genes associated to CRISPR-Cas genomes (GO Biological Process and Molecular function).

**Suppl. Fig. S4.**
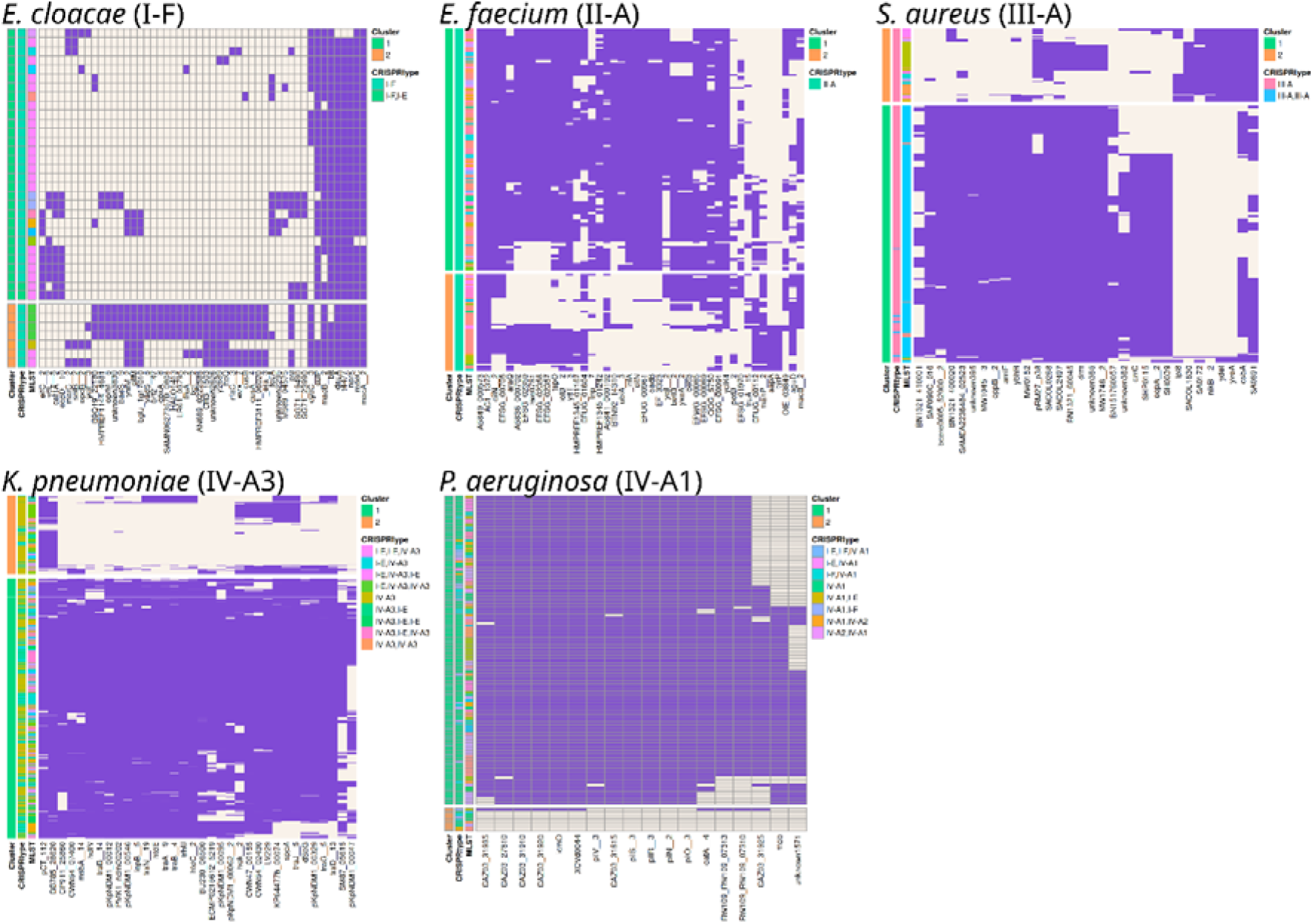
Clusters of genomes with CRISPR-Cas types II, III and IV systems, together with the I-F of *E. cloacae* by the sets of membrane protein encoding genes that they present. The purple color indicates presence of the gene of the X-axis in the corresponding genomes of the Y-axis. At the left side of each heatmap the cluster number is shown, together with the CRISPR-Cas type or combination of them and the MLST group.

**Suppl. Fig. S5.**
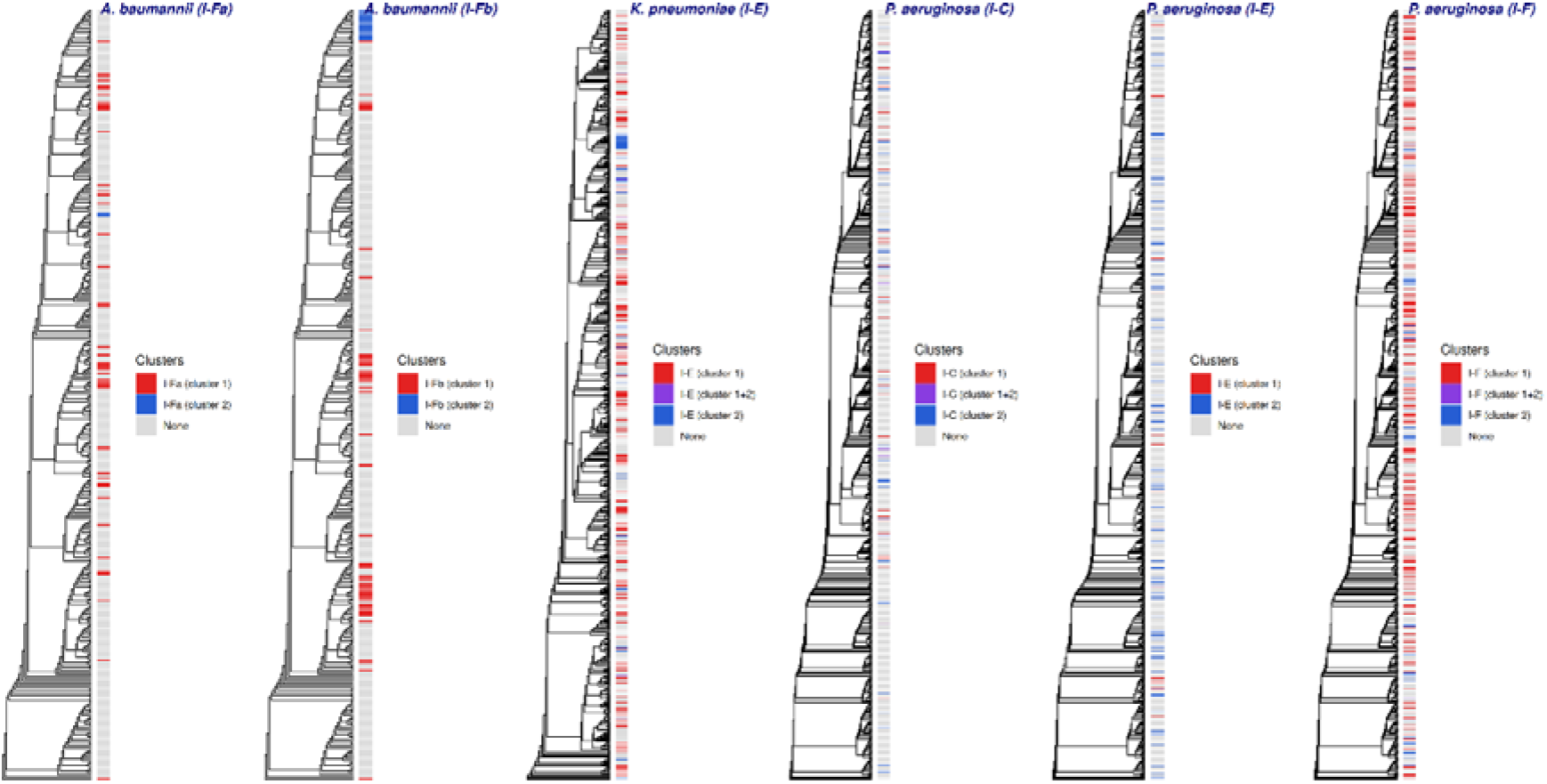
Molecular phylogenies of MLST groups separated by species and by CRISPR-Cas subtype. Nodes including genomes with CRISPR-Cas systems belonging to the clusters in Figure 4 are highlighted in different colors: cluster 1 (red color), cluster 2 (blue color) and both clusters (purple color).

**Suppl. Fig. S6.**
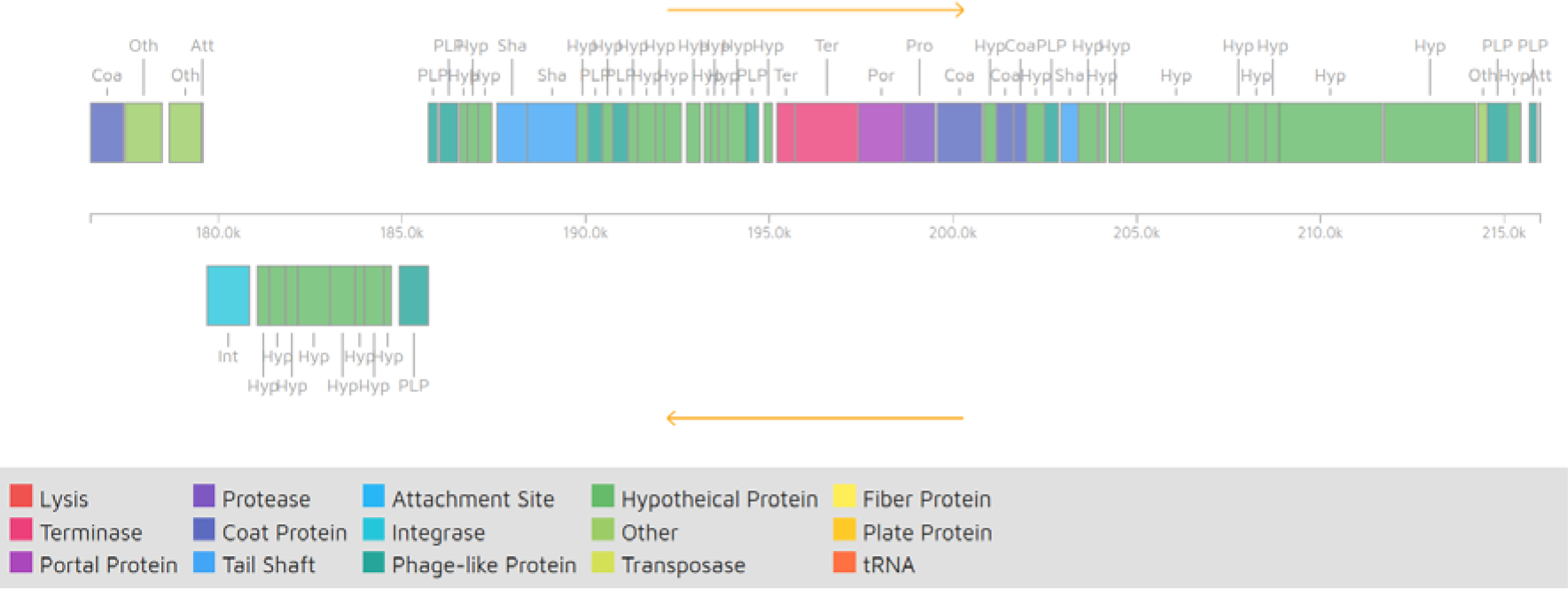
Phage with gene *unknown5433* integrated near the tRNA-Arg in *A. baumannii* (figure obtained with Phaster).

**Suppl. Table S1. Distribution of CRISPR-Cas systems and spacers among all species**.

**Suppl. Table S2. Genes associated to CRISPR-Cas genomes**. Each worksheet represents the inference results of 20 iterations of a random forest for a species and the respective CRISPR-Cas systems. The first worksheet has a header explaining the columns as follows: genes (holds the encoded gene names); known_genes_retained (contains the count of how often the gene was in the top 30 most important features of the trained random forest); log2fc_gene and diff_gene, which are the log2 fold change and the simple difference of means of the samples with and without a CRISPR-Cas system, respectively.

**Suppl. Table S3. Genomes used for each species with the MLST to which they belong, and the CRISPR-Cas systems found in them**. Incomplete CRISPR-Cas systems are labeled as “ambiguous”. The *A. baumannii* I-Fb clusters have been added, along with the metadata colors used in Fig. 6d.

## References

1. Antimicrobial Resistance Collaborators, Global burden of bacterial antimicrobial resistance in 2019: a systematic analysis. Lancet 399, 629–655 (2022).

2. E. Tacconelli, et al., Discovery, research, and development of new antibiotics: the WHO priority list of antibiotic-resistant bacteria and tuberculosis. Lancet Infect Dis 18, 318–327 (2018).

3. R. Tommasi, D. G. Brown, G. K. Walkup, J. I. Manchester, A. A. Miller, ESKAPEing the labyrinth of antibacterial discovery. Nat Rev Drug Discov 14, 529–542 (2015).

4. E. Avershina, V. Shapovalova, G. Shipulin, Fighting Antibiotic Resistance in Hospital-Acquired Infections: Current State and Emerging Technologies in Disease Prevention, Diagnostics and Therapy. Front Microbiol 12, 707330 (2021).

5. H. Nikaido, Molecular basis of bacterial outer membrane permeability revisited. Microbiol Mol Biol Rev 67, 593–656 (2003).

6. R. M. Dedrick, et al., Phage Therapy of Mycobacterium Infections: Compassionate-use of Phages in Twenty Patients with Drug-Resistant Mycobacterial Disease. Clin Infect Dis, ciac453 (2022).

7. F. Tesson, et al., Systematic and quantitative view of the antiviral arsenal of prokaryotes. Nat Commun 13, 2561 (2022).

8. L. A. Marraffini, E. J. Sontheimer, CRISPR interference: RNA-directed adaptive immunity in bacteria and archaea. Nat Rev Genet 11, 181–190 (2010).

9. R. M. Wheatley, R. C. MacLean, CRISPR-Cas systems restrict horizontal gene transfer in Pseudomonas aeruginosa. ISME J 15, 1420–1433 (2021).

10. K. Mortensen, T. J. Lam, Y. Ye, Comparison of CRISPR-Cas Immune Systems in Healthcare-Related Pathogens. Front Microbiol 12, 758782 (2021).

11. S. A. Shmakov, et al., The CRISPR Spacer Space Is Dominated by Sequences from Species-Specific Mobilomes. MBio 8 (2017).

12. G. E. Heussler, et al., Clustered Regularly Interspaced Short Palindromic Repeat-Dependent, Biofilm-Specific Death of Pseudomonas aeruginosa Mediated by Increased Expression of Phage-Related Genes. mBio 6, e00129–00115 (2015).

13. B. Tang, et al., Deletion of cas3 gene in Streptococcus mutans affects biofilm formation and increases fluoride sensitivity. Arch. Oral Biol. 99, 190–197 (2019).

14. K. C. Cady, G. A. O’Toole, Non-identity-mediated CRISPR-bacteriophage interaction mediated via the Csy and Cas3 proteins. J Bacteriol 193, 3433–3445 (2011).

15. L. Medina-Aparicio, et al., The CRISPR-Cas System Is Involved in OmpR Genetic Regulation for Outer Membrane Protein Synthesis in Salmonella Typhi. Front Microbiol 12, 657404 (2021).

16. T. R. Sampson, S. D. Saroj, A. C. Llewellyn, Y.-L. Tzeng, D. S. Weiss, A CRISPR/Cas system mediates bacterial innate immune evasion and virulence. Nature 497, 254–257 (2013).

17. M. A. B. Shabbir, et al., The Involvement of the Cas9 Gene in Virulence of Campylobacter jejuni. Front Cell Infect Microbiol 8, 285 (2018).

18. J. Solbiati, A. Duran-Pinedo, F. Godoy Rocha, F. C. Gibson, J. Frias-Lopez, Virulence of the Pathogen Porphyromonas gingivalis Is Controlled by the CRISPR-Cas Protein Cas3. mSystems 5, e00852–20 (2020).

19. E. L. Mangas, et al., Pangenome of Acinetobacter baumannii uncovers two groups of genomes, one of them with genes involved in CRISPR/Cas defence systems associated with the absence of plasmids and exclusive genes for biofilm formation. Microb Genom 5 (2019).

20. Y. Kim, C. Gu, H. U. Kim, S. Y. Lee, Current status of pan-genome analysis for pathogenic bacteria. Curr Opin Biotechnol 63, 54–62 (2020).

21. R. J. Hall, F. J. Whelan, J. O. McInerney, Y. Ou, M. R. Domingo-Sananes, Horizontal Gene Transfer as a Source of Conflict and Cooperation in Prokaryotes. Front Microbiol 11, 1569 (2020).

22. P. A. Kitts, et al., Assembly: a resource for assembled genomes at NCBI. Nucleic Acids Res. 44, D73–80 (2016).

23. T. Seemann, Prokka: rapid prokaryotic genome annotation. Bioinformatics 30, 2068–2069 (2014).

24. A. J. Page, et al., Roary: rapid large-scale prokaryote pan genome analysis. Bioinformatics 31, 3691–3693 (2015).

25. C. S. Casimiro-Soriguer, A. Muñoz-Mérida, A. J. Pérez-Pulido, Sma3s: A universal tool for easy functional annotation of proteomes and transcriptomes. Proteomics 17 (2017).

26. J. Russel, R. Pinilla-Redondo, D. Mayo-Muñoz, S. A. Shah, S. J. Sørensen, CRISPRCasTyper: Automated Identification, Annotation, and Classification of CRISPR-Cas Loci. CRISPR J 3, 462–469 (2020).

27. D. Couvin, et al., CRISPRCasFinder, an update of CRISRFinder, includes a portable version, enhanced performance and integrates search for Cas proteins. Nucleic Acids Res. 46, W246–W251 (2018).

28. M. Feldgarden, et al., AMRFinderPlus and the Reference Gene Catalog facilitate examination of the genomic links among antimicrobial resistance, stress response, and virulence. Sci Rep 11, 12728 (2021).

29. C. Camacho, et al., BLAST+: architecture and applications. BMC Bioinformatics 10, 421 (2009).

30. B. Liu, D. Zheng, Q. Jin, L. Chen, J. Yang, VFDB 2019: a comparative pathogenomic platform with an interactive web interface. Nucleic Acids Res 47, D687–D692 (2019).

31. K. D. Tsirigos, P. G. Bagos, S. J. Hamodrakas, OMPdb: a database of {beta}-barrel outer membrane proteins from Gram-negative bacteria. Nucleic Acids Res 39, D324–331 (2011).

32. V. Galata, T. Fehlmann, C. Backes, A. Keller, PLSDB: a resource of complete bacterial plasmids. Nucleic Acids Res 47, D195–D202 (2019).

33. S. Roux, et al., IMG/VR v3: an integrated ecological and evolutionary framework for interrogating genomes of uncultivated viruses. Nucleic Acids Res 49, D764–D775 (2021).

34. D. Arndt, et al., PHASTER: a better, faster version of the PHAST phage search tool. Nucleic Acids Res 44, W16–21 (2016).

35. A. Rubio, et al., CRISPR sequences are sometimes erroneously translated and can contaminate public databases with spurious proteins containing spaced repeats. Database (Oxford) 2020 (2020).

36. F. Pedregosa, et al., Scikit-learn: Machine Learning in Python. Journal of Machine Learning Research 12, 2825–2830 (2011).

37. A. Alexa, J. Rahnenführer, T. Lengauer, Improved scoring of functional groups from gene expression data by decorrelating GO graph structure. Bioinformatics 22, 1600–1607 (2006).

38. K. A. Jolley, J. E. Bray, M. C. J. Maiden, Open-access bacterial population genomics: BIGSdb software, the PubMLST.org website and their applications. Wellcome Open Res 3, 124 (2018).

39. K. Katoh, D. M. Standley, MAFFT multiple sequence alignment software version 7: improvements in performance and usability. Mol. Biol. Evol. 30, 772–780 (2013).

40. A. Stamatakis, RAxML version 8: a tool for phylogenetic analysis and post-analysis of large phylogenies. Bioinformatics 30, 1312–1313 (2014).

41. S. Kalyaanamoorthy, B. Q. Minh, T. K. F. Wong, A. von Haeseler, L. S. Jermiin, ModelFinder: fast model selection for accurate phylogenetic estimates. Nat Methods 14, 587–589 (2017).

42. P. Peterson, F2PY: a tool for connecting Fortran and Python programs. Int. J. Comput. Sci. Eng. 4, 296–305 (2009).

43. M. L. Waskom, seaborn: statistical data visualization. Journal of Open Source Software 6, 3021 (2021).

44. J. D. Hunter, Matplotlib: A 2D Graphics Environment. Computing in Science Engineering 9, 90–95 (2007).

45. E. Pfeifer, J. A. Moura de Sousa, M. Touchon, E. P. C. Rocha, Bacteria have numerous distinctive groups of phage-plasmids with conserved phage and variable plasmid gene repertoires. Nucleic Acids Res 49, 2655–2673 (2021).

46. A. Moya-Beltrán, et al., Evolution of Type IV CRISPR-Cas Systems: Insights from CRISPR Loci in Integrative Conjugative Elements of Acidithiobacillia. The CRISPR Journal 4, 656–672 (2021).

47. J. Breisch, B. Averhoff, Identification of osmo-dependent and osmo-independent betaine-choline-carnitine transporters in Acinetobacter baumannii: role in osmostress protection and metabolic adaptation. Environ Microbiol 22, 2724–2735 (2020).

48. L. Moynié, et al., Preacinetobactin not acinetobactin is essential for iron uptake by the BauA transporter of the pathogen Acinetobacter baumannii. Elife 7, e42270 (2018).

49. J. Bertozzi Silva, Z. Storms, D. Sauvageau, Host receptors for bacteriophage adsorption. FEMS Microbiol Lett 363, fnw002 (2016).

50. P. Bernal, L. P. Allsopp, A. Filloux, M. A. Llamas, The Pseudomonas putida T6SS is a plant warden against phytopathogens. ISME J 11, 972–987 (2017).

51. K. D. Mlaga, et al., Extensive Comparative Genomic Analysis of Enterococcus faecalis and Enterococcus faecium Reveals a Direct Association between the Absence of CRISPR-Cas Systems, the Presence of Anti-Endonuclease (ardA) and the Acquisition of Vancomycin Resistance in E. faecium. Microorganisms 9, 1118 (2021).

52. E. Pursey, T. Dimitriu, F. L. Paganelli, E. R. Westra, S. van Houte, CRISPR-Cas is associated with fewer antibiotic resistance genes in bacterial pathogens. Philos Trans R Soc Lond B Biol Sci 377, 20200464 (2022).

53. J. McGinn, L. A. Marraffini, Molecular mechanisms of CRISPR-Cas spacer acquisition. Nat Rev Microbiol 17, 7–12 (2019).

54. S. A. Shmakov, Y. I. Wolf, E. Savitskaya, K. V. Severinov, E. V. Koonin, Mapping CRISPR spaceromes reveals vast host-specific viromes of prokaryotes. Commun Biol 3, 321 (2020).

55. U. Gophna, et al., No evidence of inhibition of horizontal gene transfer by CRISPR-Cas on evolutionary timescales. ISME J 9, 2021–2027 (2015).

56. G. Faure, et al., CRISPR-Cas in mobile genetic elements: counter-defence and beyond. Nat. Rev. Microbiol. (2019) https://doi.org/10.1038/s41579-019-0204-7.

57. R. Perez-Rodriguez, et al., Envelope stress is a trigger of CRISPR RNA-mediated DNA silencing in Escherichia coli. Mol Microbiol 79, 584–599 (2011).

58. S. A. Shmakov, K. S. Makarova, Y. I. Wolf, K. V. Severinov, E. V. Koonin, Systematic prediction of genes functionally linked to CRISPR-Cas systems by gene neighborhood analysis. Proc. Natl. Acad. Sci. U.S.A. 115, E5307–E5316 (2018).

59. S. A. Shah, et al., Comprehensive search for accessory proteins encoded with archaeal and bacterial type III CRISPR-cas gene cassettes reveals 39 new cas gene families. RNA Biol 16, 530–542 (2019).

60. L.-C. Fortier, O. Sekulovic, Importance of prophages to evolution and virulence of bacterial pathogens. Virulence 4, 354–365 (2013).

61. A. Stern, L. Keren, O. Wurtzel, G. Amitai, R. Sorek, Self-targeting by CRISPR: gene regulation or autoimmunity? Trends Genet. 26, 335–340 (2010).

62. S. Govindarajan, A. Borges, S. Karambelkar, J. Bondy-Denomy, Distinct Subcellular Localization of a Type I CRISPR Complex and the Cas3 Nuclease in Bacteria. J Bacteriol 204, e0010522 (2022).

63. G. W. Goldberg, W. Jiang, D. Bikard, L. A. Marraffini, Conditional tolerance of temperate phages via transcription-dependent CRISPR-Cas targeting. Nature 514, 633–637 (2014).

64. N.-H. Le, V. Pinedo, J. Lopez, F. Cava, M. F. Feldman, Killing of Gram-negative and Gram-positive bacteria by a bifunctional cell wall-targeting T6SS effector. Proc Natl Acad Sci U S A 118, e2106555118 (2021).

65. L. Letellier, P. Boulanger, L. Plançon, P. Jacquot, M. Santamaria, Main features on tailed phage, host recognition and DNA uptake. Front Biosci 9, 1228–1339 (2004).

66. L. Cui, et al., CRISPR-cas3 of Salmonella Upregulates Bacterial Biofilm Formation and Virulence to Host Cells by Targeting Quorum-Sensing Systems. Pathogens 9, E53 (2020).

67. A. D. Maharajan, E. Hjerde, H. Hansen, N. P. Willassen, Quorum Sensing Controls the CRISPR and Type VI Secretion Systems in Aliivibrio wodanis 06/09/139. Front Vet Sci 9, 799414 (2022).

