## Supplementary figures and images for "Reducing CRISPR dark matter reveals a strong association between the bacterial membranome and CRISPR-Cas systems"

### Suppl. Fig. S1

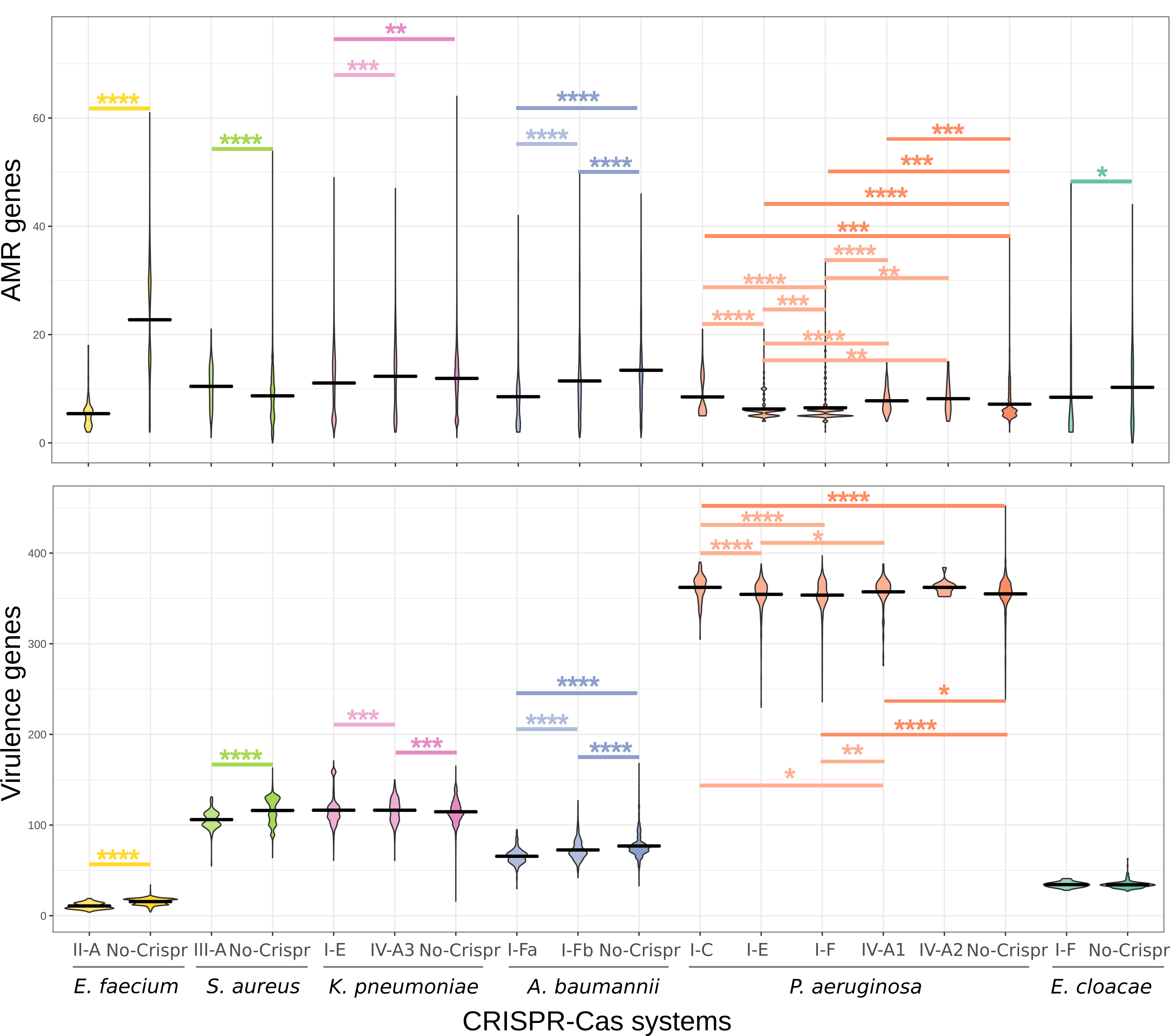

### Suppl. Fig. S2

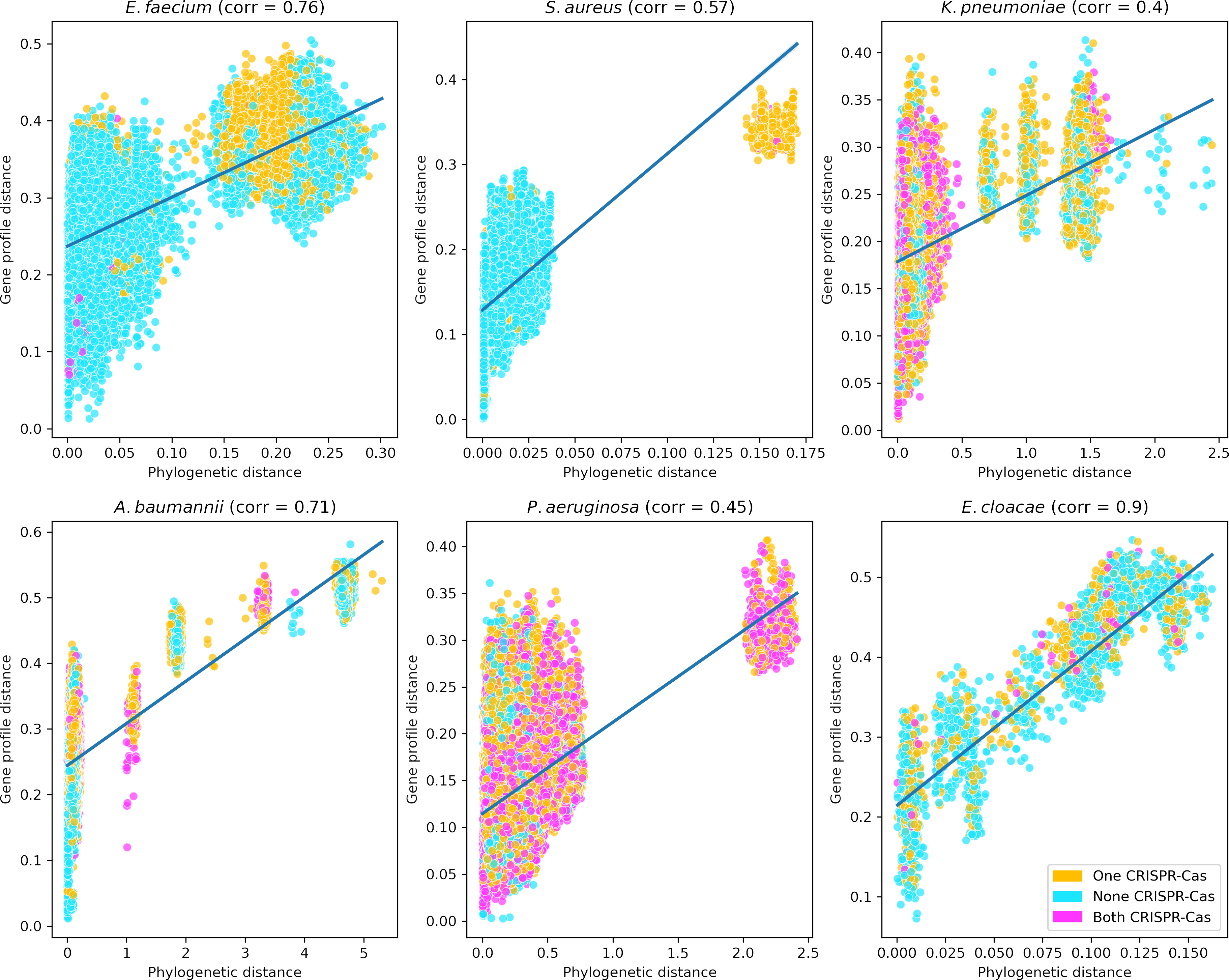

### Suppl. Fig. S3

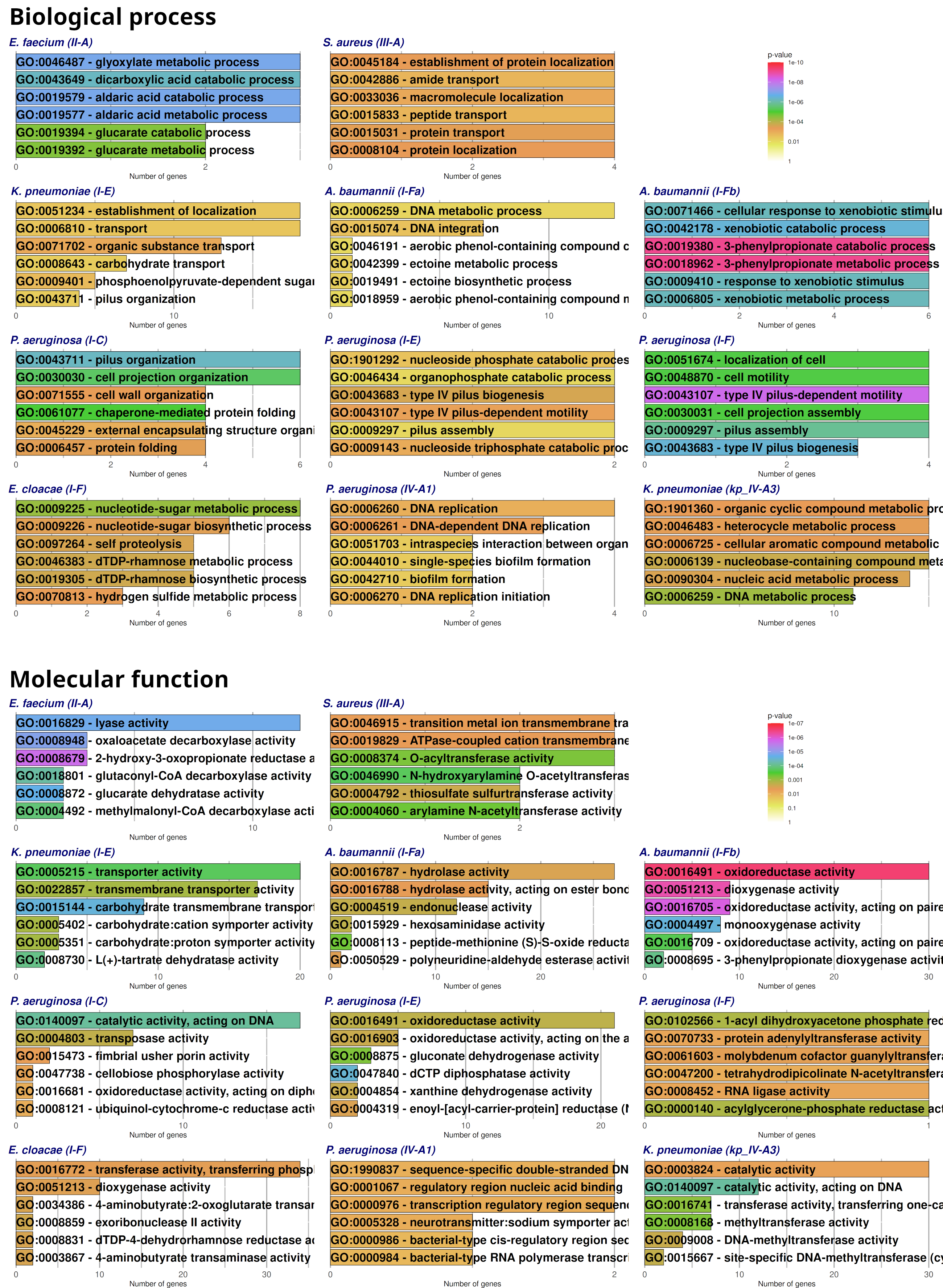

### Suppl. Fig. S4

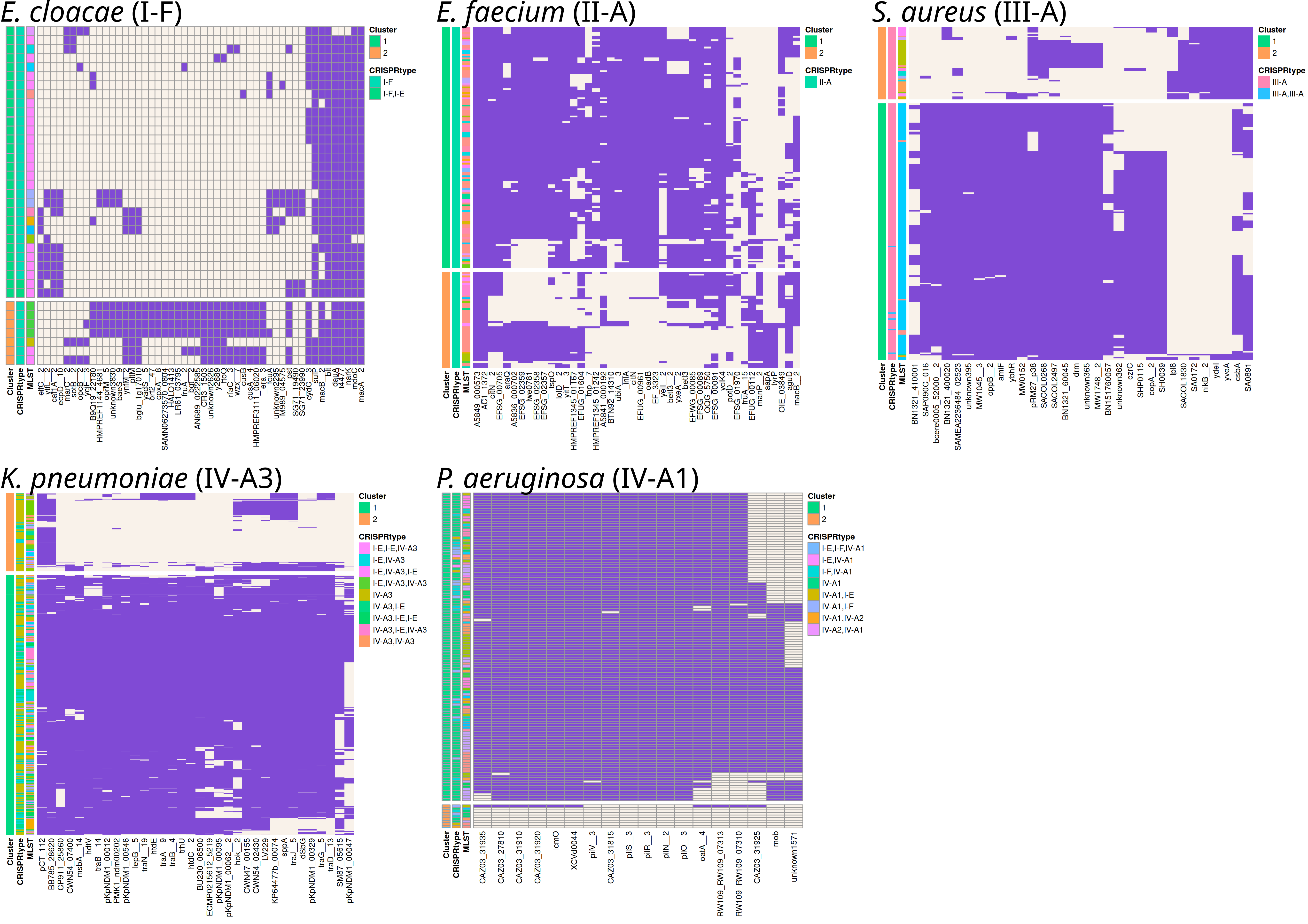

### Suppl. Fig. S5

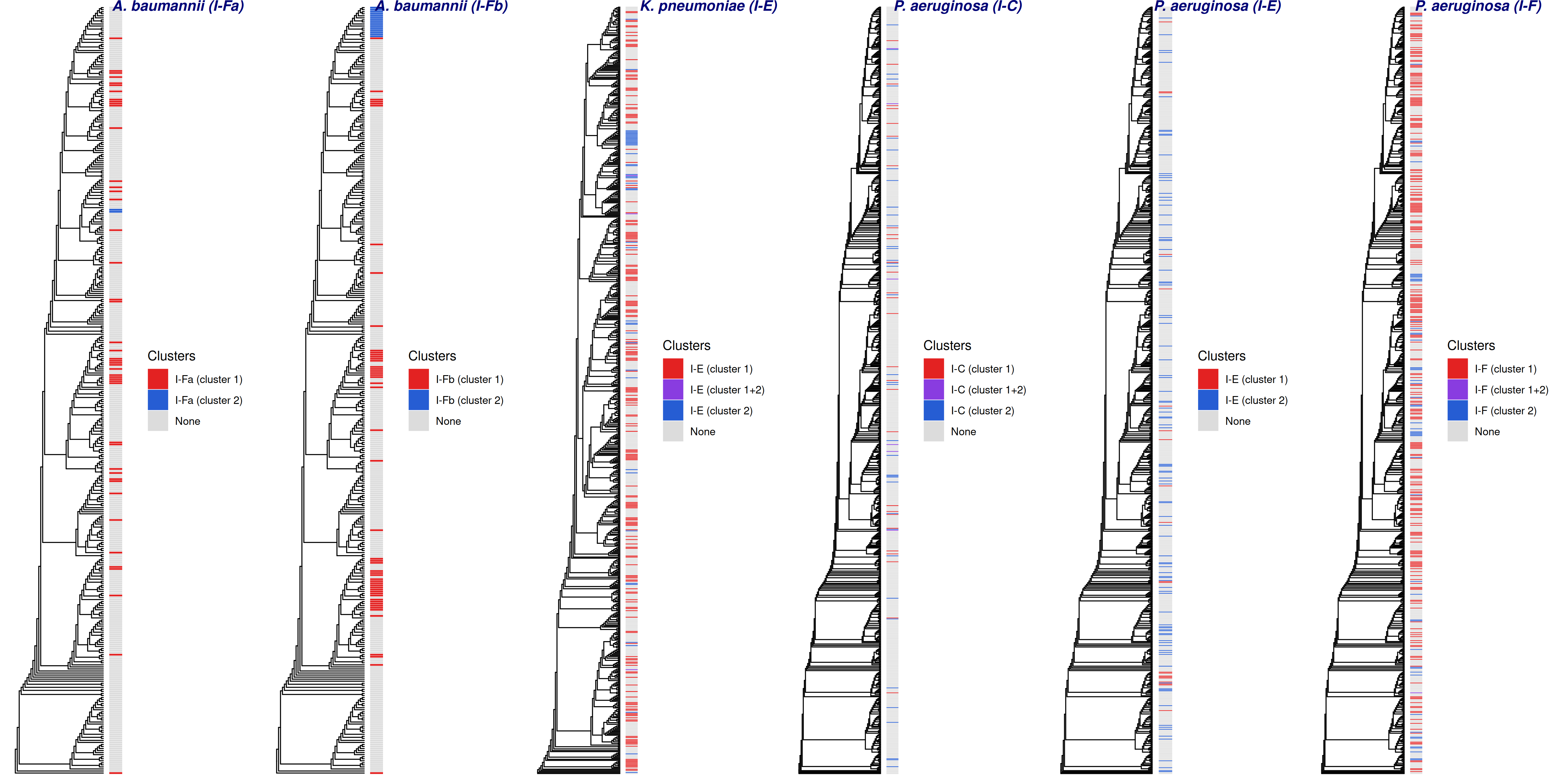

### Suppl. Fig. S6

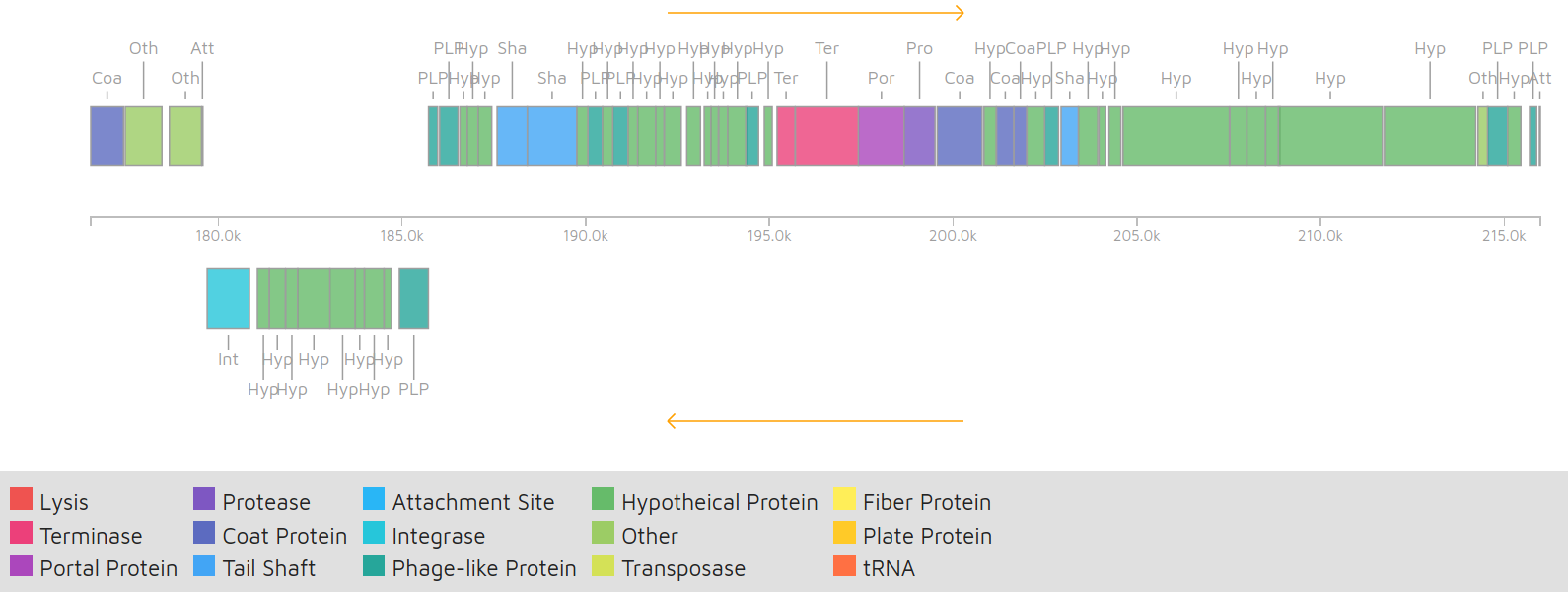
